# PERK Signaling Maintains Hematopoietic Stem Cell Pool Integrity under Endoplasmic Reticulum Stress by Promoting Proliferation

**DOI:** 10.1101/2024.12.13.628451

**Authors:** Manxi Zheng, Qinlu Peng, Erin M Kropp, Zhejuan Shen, Suxuan Liu, Zhengyou Yin, Sho Matono, Takao Iwawaki, Xiang Wang, Ken Inoki, Yang Mei, Qing Li, Lu Liu

## Abstract

The integrity of the hematopoietic stem cell (HSC) pool relies on efficient long-term self-renewal and the timely removal of damaged or differentiation-prone HSCs. Previous studies have demonstrated the PERK branch of the unfolded protein response (UPR) drives specific programmed cell death programs to maintain HSC pool integrity in response to ER stress. However, the role of PERK in regulating HSC fate *in vivo* remains unclear. Here, we demonstrate that PERK is dispensable for normal hematopoiesis and HSC self-renewal under steady-state conditions. In contrast, PERK is activated to promote HSC proliferation and depletion in response to ER stress induced by the inactivation of ER-associated degradation (ERAD), via the knockout of key components of ERAD Sel1L or Hrd1. Inhibition of PERK, either through genetic knockout or knock-in of a point mutation that eliminates PERK kinase activity, significantly restores the HSC defects induced by Sel1L or Hrd1 knockout. Mechanistic studies reveal that ERAD deficiency does not lead to HSC death or ROS accumulation. Instead, PERK promotes the activation of mTOR signaling and drives abnormal proliferation of HSCs, impairing their self-renewal potential. This process removes stressed HSCs, thereby maintaining HSC pool integrity. Our study uncovers a PERK-centered strategy employed by HSCs to preserve their pool integrity independently of apoptosis.

**Key points:**

1. PERK is not required for steady-state hematopoiesis but preserves hematopoietic stem cell pool integrity in response to increased ER stress.
2. Under ER stress induced by ERAD deficiency, PERK is activated to promote mTOR signaling and HSC hyper-proliferation, depleting damaged HSCs.

## Introduction

Hematopoietic stem cells (HSCs) generate all the hematopoietic progenitors, which subsequently differentiate into lineage cells^1–3^. To maintain life-time hematopoiesis, the integrity of HSC pool must be tightly maintained^4,5^. HSCs are primarily maintained in a quiescent state to sustain long-term self-renewal ^6–10^. Signals that trigger proliferation and activation of HSCs frequently lead to a significant decline in self-renewal potential and promote differentiation^11^. Similarly, under stress conditions or in the presence of extrinsic or intrinsic defects, HSCs exit from the quiescent HSC pool by hyper-proliferation and differentiation ^8,11^. Hence, proliferation and differentiation could eliminate damaged HSCs from the quiescent pool to maintain HSC integrity independent of cell death.

Protein homeostasis is crucial for HSC functions and must be precisely regulated by a network of protein quality control (PQC) mechanisms. Endoplasmic Reticulum (ER) associated degradation (ERAD) and unfolded protein response (UPR)^12–14^ govern protein homoestasis in ER, which is responsible for up to 30% of total protein synthesis^15^. As ER is susceptable to inherited protein misfolding and stress conditions, in response to ER stress, ERAD is transcriptional activated by UPR to promote degradation of misfolded proteins ^16^. Sel1l-Hrd1 ERAD complex has been proved to be essential for maintaining HSC quiescence to sustain long-term hematopoiesis. The loss of Sel1l results in a complete loss in self-renewal potential of HSCs by increasing proliferation without inducing apoptosis in HSCs^13,17^. While these studies established the role of ERAD in maintaining HSC quiescence and self-renewal, the signaling mechanism by which this occurs and how ER stress response pathways respond to impaired proteostasis remain to be investigated.

The UPR is activiated upon increased concentration of unfolded or misfolded proteins in the ER^18^. This pathway is transduced by three sensors: inositol-requiring enzyme 1α (IRE1α), protein kinase R-like endoplasmic reticulum kinase (PERK), and activating transcription factor 6 (ATF6)^19^. While early responses can mitigate ER stress and protein burden through ATF6 and IRE1α branches, persistent ER stress has been associated with an induction of apoptosis in a PERK-dependent manner to maintain the integrity of the cells^20^. In this process, PERK phosphorylates eIF2α to promotes the translation of a set of “stress” related genes including ATF4^21^. ATF4 further induces CHOP expression to promote apoptosis^19^. The impact of UPR activation is highly cell-type and context-dependent. PERK was previously shown to promote apoptosis in HSCs under extreme ER stress induced by tunicamycin or thapsigargin *in vitro*. The role of PERK in HSCs *in vivo*, either in steady-state or under ER stress remains to be investigated.

In this study, we evaluate the role of PERK in HSC quiescence, hematopoiesis, and response to altered ER proteostasis induced by ERAD deficiency. Our findings suggest that PERK is dispensable for hematopoiesis, HSC pool size and HSC function in steady state. However, under increased ER stress induced by ERAD deficiency, via Sel1l or Hrd1 knockout, PERK is activated to promote HSC hyperproliferation and depletion, and PERK knockout mitigates the HSC impairments induced by Sel1l or Hrd1 knockout. Mechanistic studies revealed that the effects of PERK are mediated by its kinase activity, and the knock-in of a mutant PERK that lacks the kinase activity blocked the effects of PERK in HSCs. Our studies, therefore, indicate an essential role of PERK in guarding the integrity of HSCs by eliminating damaged HSCs after ER stress through hyper-proliferation.

## Methods

### Mice

Mice were housed in the Unit for Laboratory Animal Medicine (ULAM) at the University of Michigan and the Laboratory Animal Center of Hunan University. All procedures involving mouse use and care were conducted in accordance with the ethical regulations and policies established by the University of Michigan Institutional Animal Care and Use Committee (IACUC) and Hunan University IACUC. All animal protocols were reviewed and approved by the respective IACUCs. The mice used in this study are listed in **Supplementary Table 1** and were maintained on a C57BL background. Other and detailed description of methods are attached as **Supplementary methods** due to the words limits.

## Results

### PERK is dispensable for steady stage hematopoiesis, HSC proliferation and reconstitution

To determine the role of PERK signaling in HSCs and hematopoiesis, we crossed *Mx1-cre^+^* mice^22^ with conditional PERK knockout mice^23^ to generate *Mx1-cre^+^; PERK ^fl/fl^* mice. After poly-inosine-poly-cytosine (pIpC; 0.5 µg/gram body weight every other day for six doses) induced deletion of PERK in hematopoietic tissues. These mice were then compared to sex- and age-matched *Mx1-cre^-^; PERK ^fl/fl^* mice (control, +/+). Two weeks after pIpC injection, PERK knockout (KO) was confirmed by Western Blot (**Supplementary Figure 1A**). Homozygous PERK KO mice exhibited similar body weight, bone marrow, and spleen cellularity, as well as comparable spleen and thymus weights and cellularity to control mice (**Figure 1A-F** and **Supplementary Figure 1B-C**).

**Figure 1.**
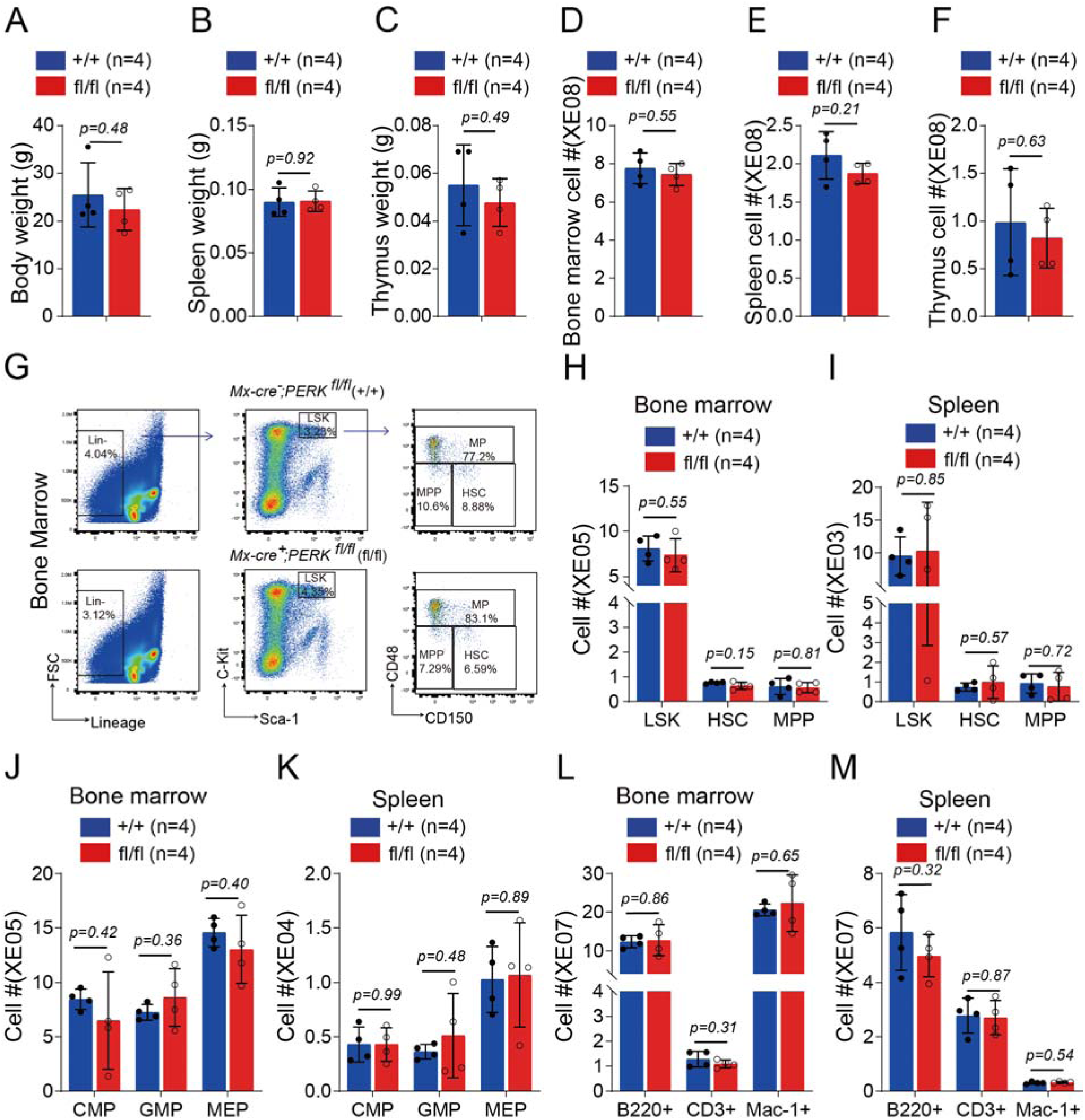
PERK is dispensable for steady state hematopoiesis. 6*^-^*8 weeks old *Mx1-cre^-^; PERK^fl/fl^* (+/+) and *Mx1-cre^+^; PERK^fl/fl^* (fl/fl) mice were injected with pIpC every other day for a total of 6 doses. Two weeks after pIpC injection, body weight **(A)**, spleen and thymus weight (**B-C)**, cellularity of bone marrow, spleen and thymus (**D*-*F**), numbers of HSPC (**G*-*I**), lineage restricted progenitors (**J-K**) and mature blood cells (**L-M**) were analyzed. Data represent mean±s.d. Each replicate represents a single mouse from an independent experiment.

PERK KO displayed comparable numbers and frequencies of SLAM^24^ HSCs, multipotent progenitors (MPP; CD150*^-^*CD48*^-^*LSK)(**Supplementary Figure 1D-E** and **Figure 1G-I**), common myeloid progenitor (CMP; Lineage*^-^*Sca1*^-^*cKit*^+^*CD16/32*^+^*CD34^-^), granulocyte-macrophage progenitor (GMP; Lineage*^-^*Sca1*^-^*cKit*^+^* CD16/32*^+^*CD34*^+^*) and megakaryocyte-erythroid progenitor (MEP; Lineage*^-^*Sca1*^-^*cKit*^+^* CD16/32^-^CD34^-^)(**Supplementary Figure 1F-G** and **Figure 1J-K**), mature myeloid, B, and T lineage cells(**Figure 1L-M** and **Supplementary Figure 1H-I**) in the spleen and bone marrow comparing to control mice. These findings suggest that PERK deletion does not affect hematopoietic stem cells and hematopoiesis under steady-state conditions.

The phenotype of *Mx1-cre^-^*driven PERK KO was confirmed by crossing *Vav1-cre*^25^ mice with *PERK ^fl/fl^* mice. *Vav1-cre^+^; PERK ^fl/fl^* mice and control mice demonstrated similar steady stage hematopoisis (**Supplementary Figure 2A-R**). These results confirmed that PERK deletion does not impact hematopoietic populations under steady-state conditions.

To evaluate the reconstitution potential of HSCs, we transplanted 100 FACS-purified HSCs from 6-8 week-old CD45.2*^+^ Vav1-cre^+^*; *PERK ^fl/fl^* or control mice, along with 5x10^5^ CD45.1*^+^* wild-type bone marrow cells, into lethally irradiated CD45.1*^+^* wild-type recipients. The transplants of PERK KO and control HSCs did not exhibit a significant difference in donor reconstitution in total CD45*^+^*, myeloid (Mac1*^+^*), B (B220*^+^*), and T (CD3*^+^*) cells 4-16 weeks after transplantation (**Figure 2A**). Analysis of the bone marrow at 16 weeks after transplantation showed that donor reconstitution in HSCs and all other hematopoietic populations was comparable in transplant recipients between PERK KO and controls (**Figure 2B**), confirming no significant change in HSC reconstitution. These results collectively demonstrate that UPR PERK signaling is dispensable for HSC reconstitution and hematopoiesis at a steady state.

**Figure 2.**
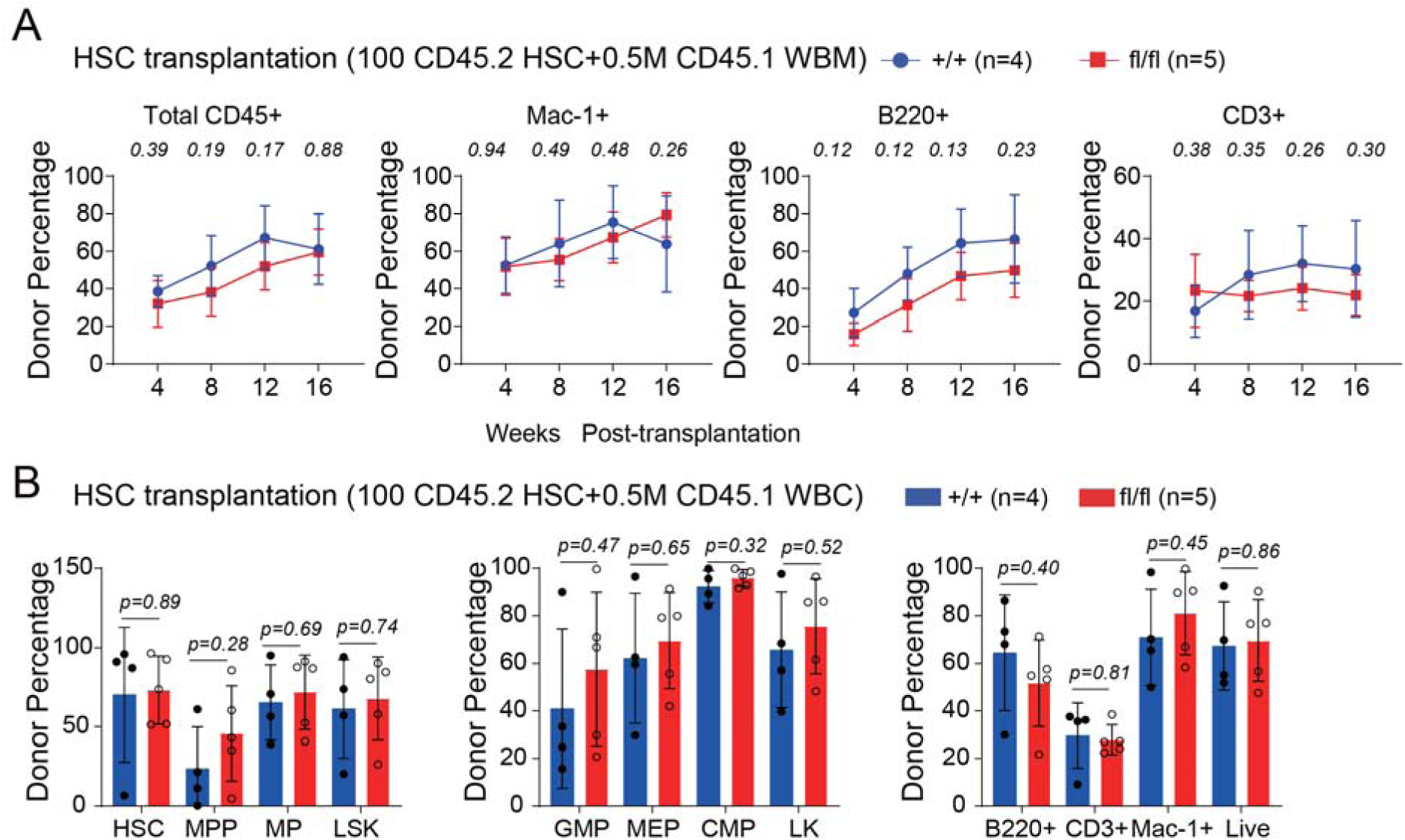
PERK is not required for HSC reconstitution in transplants. 100 FACS*-*purified HSCs from CD45.2 *Vav-cre^+^; PERK^fl/fl^* (fl/fl) or control (*+*/*+*) mice were transplanted into irradiated receipt mice along with 0.5 million CD45.1 whole bone marrow competitors. **A).** The contribution of CD45.2 cells in total CD45*^+^*, myeloid (Mac*-*1*^+^*), B (B220*^+^*) and T (CD3*^+^*) cells in peripheral blood was analyze every 4 weeks for 16 weeks. **B).** The contribution of CD45.2 cells in HSC, progenitors and mature lineage cells in the bone marrow was analyzed at the end of 16 weeks. Data represent mean±s.d. Each replicate represents a single mouse. Two*^-^*sided student t*^-^*test was used for statistical analysis. P values are included on the top of the graphs.

### PERK was activated in response to non-apoptotic ER stress in HSCs

Although PERK has been shown to promote cell death of HSCs *in vitro* in response to severe ER stress triggered by tunicamycin or thapsigargin^20^, the role of PERK in response to ER stress *in vivo* remains unclear. ERAD deficiency leads to the accumulation of misfolded proteins in the ER, which subsequently induces ER stress and activates the UPR^26^. Although persistent ER stress can result in cell death, our previous work demonstrated that impaired ERAD activity decreases the HSC pool size without increasing apoptosis^13^. To investigate whether HSCs experience ER stress when ERAD activity is inhibited, we purified HSCs from 6-8-week-old *Mx1-cre^+^*; *Sel1L ^fl/fl^* mice (2 weeks after pIpC) and evaluated UPR by examining key targets of the UPR pathways via RT-qPCR. Expression levels of Grp78, Xbp1s, Erdj4, and Chop were significantly elevated in Sel1L KO HSCs compared to control HSCs (**Figure 3A**), indicating transcriptional activation of UPR targets. To confirm IRE1α activation, we crossed ERAI mice, a transgenic reporter mouse strain that detects Xbp1 splicing with Venus fluorescent expression ^27^, with *Mx1-cre^+^*; *Sel1L ^fl/fl^*mice and observed substantially higher levels of ERAI in all populations, including HSCs, in Sel1L KO mice compared to controls (**Supplementary Figure 3B**). These results suggest that the IRE1α-Xbp1 branch of UPR is activated in response to ERAD deficiency in hematopoietic cells.

**Figure 3.**
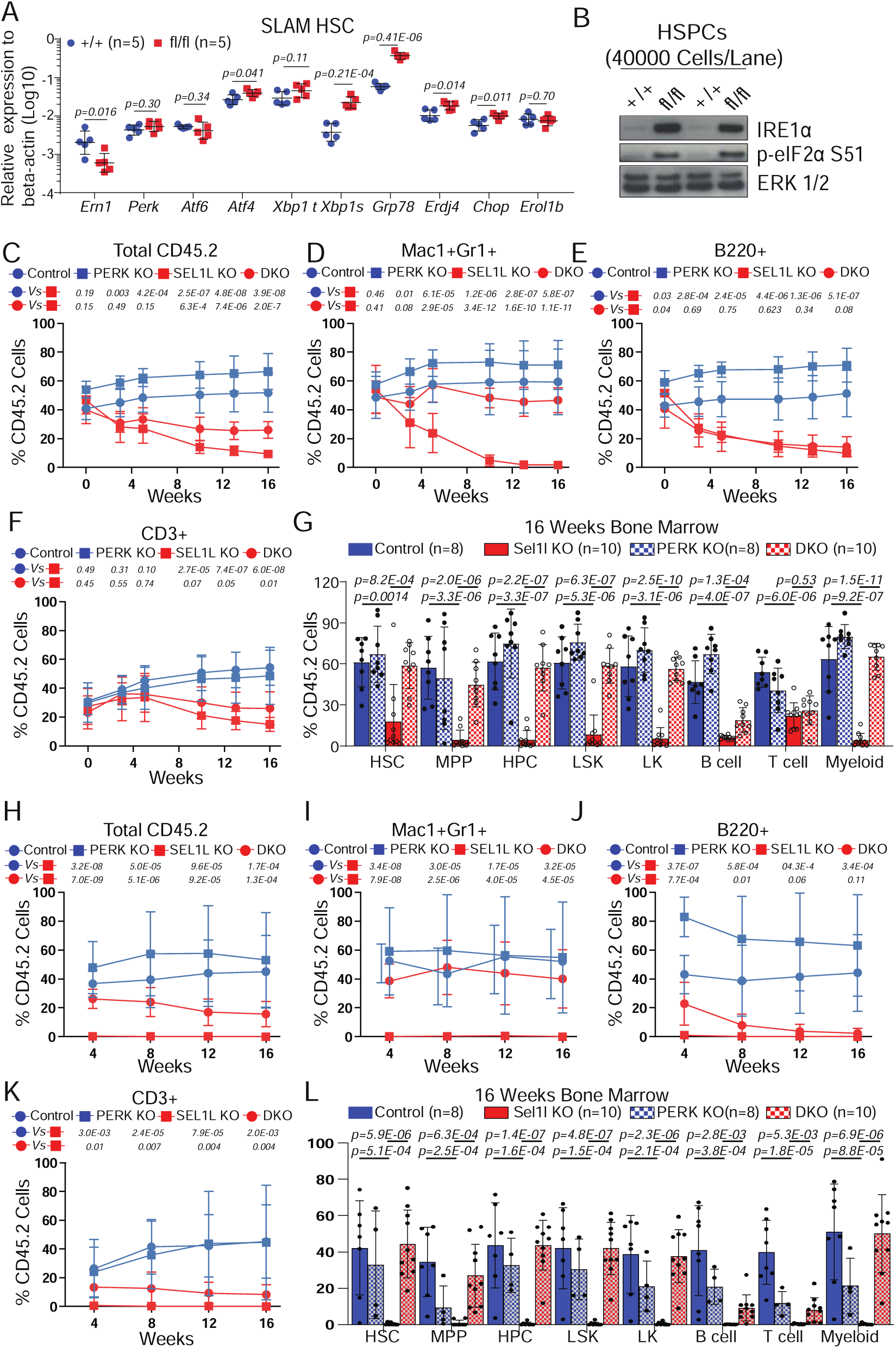
PERK is activated and mediates the impairment of HSCs under ER stress induced by Sel1L knockout. **A)**. RT-qPCR of ER stress signaling targets in purified SLAM HSCs from *Mx1-cre^+^; Sel1L^fl/fl^* (fl/fl) or control (*+*/*+*) mice two weeks after pIpC injection. **B)**. Western blot of IRE1α and phosphor-eIF2α in LSK cells (HSPCs) of *Mx1-cre^+^; Sel1L^fl/fl^* (fl/fl) and control (*+*/*+*) mice two weeks after pIpC injection. ERK level was used as loading control. **C-G)**. Chimerism maintenance analysis with 5x10^5^ CD45.2*^+^* whole bone marrow cells from *Mx1^-^cre^+^; Sel1L ^fl/fl^* (Sel1L KO)*, Mx1-cre^+^; PERK ^fl/fl^* (PERK KO), *Mx1-cre^+^*; *Sel1L ^fl/fl;^ PERK ^fl/fl^* (DKO), or control mice (without pIpC) transplanted together with 5x10^5^ CD45.1*^+^* wild type bone marrow cells into lethally irradiated CD45.1*^+^* wild*^-^*type recipients. Transplants were injected with pIpC (3 doses) 6 weeks after transplantation. The contribution of CD45.2 cells in total CD45^+^ (**C**), myeloid (Mac1+Gr1+; **D**), B (B220; **E**) and T (CD3; **F**) cells from peripheral blood, and in HSCs and other hematopoietic populations in the bone marrow (**G**) was analyzed in transplant recipients. **H-L)**. Secondary transplantation of 3 million bone marrow cells from the primary transplants (**G**). The contribution of CD45.2 cells in total CD45^+^ (**H**), myeloid (Mac1+Gr1+; **I**), B (B220; **J**) and T (CD3; **K**) cells from peripheral blood, and in HSCs and other hematopoietic populations in the bone marrow (**L**) was analyzed in the secondary transplant recipients. Data represent mean±s.d. Each replicate represents a single mouse from two independent experiments. Two-sided student t-test was used for statistical analysis unless specified.

To assess if the other two UPR branches are also activated, western blot analysis was performed to detect the activation of the PERK and ATF6. We observed increased phosphorylation of eIF2α, indicative of activated PERK (**Figure 3B**), and elevated levels of the cleaved and activated form of ATF6 (ATF6c) in Sel1L KO LSKs (**Supplementary Figure 3A**), suggesting that all three branches of UPR are activated with the loss of Sel1L in HSPCs. Consistent with increased ER stress and UPR activation, Sel1L KO HSCs exhibited increased levels of protein aggregation (**Supplementary Figure 3C**) and increased ER volume (**Supplementary Figure 3D**).

Consistent with our previous study that did not detect increased apoptosis in the Sel1L KO HSCs (measured by annexin V staining)^13^, Sel1L KO HSCs gave rise to a comparable number of colonies in semi-solid media as the control HSCs (**Supplementary Figure 3F**). In addition, the levels of caspase 3/7 (**Supplementary Figure 3G**), ROS (**Supplementary Figure 3E**), or ER stress-associated apoptosis genes *Dr5*^28^ *and Puma*^29^ (**Supplementary Figure 3H**) were similar in Sel1L KO and control HSPC. Furthermore, treatment with tunicamycin or thapsigargin induced similar levels of apoptosis in Sel1L KO (**Supplementary Figure 3I**). These results suggest that ERAD deficiency induces non-apoptotic ER stress in HSCs *in vivo*.

To confirm the effects of ERAD deficiency on HSCs with the Sel1L KO mice, we generated a second *in vivo* model of ERAD deficiency by knocking out Hrd1, an ubiquitination E3 ligase which forms complex with Sel1L and is essential for ERAD function ^30^. We generated a new Hrd1 floxed strain where exon 1-14 of Hrd1 was flanked by loxp sites (**Figure 4A**) . The *Hrd1^fl/fl^* mice were crossed to *Vav1-Cre^+^* mice to delete Hrd1 in hematopoitic cells. Western blot analysis suggested that HRD1 protein was significantly decreased in bone marrow cells (**Figure 4B**).

**Figure 4.**
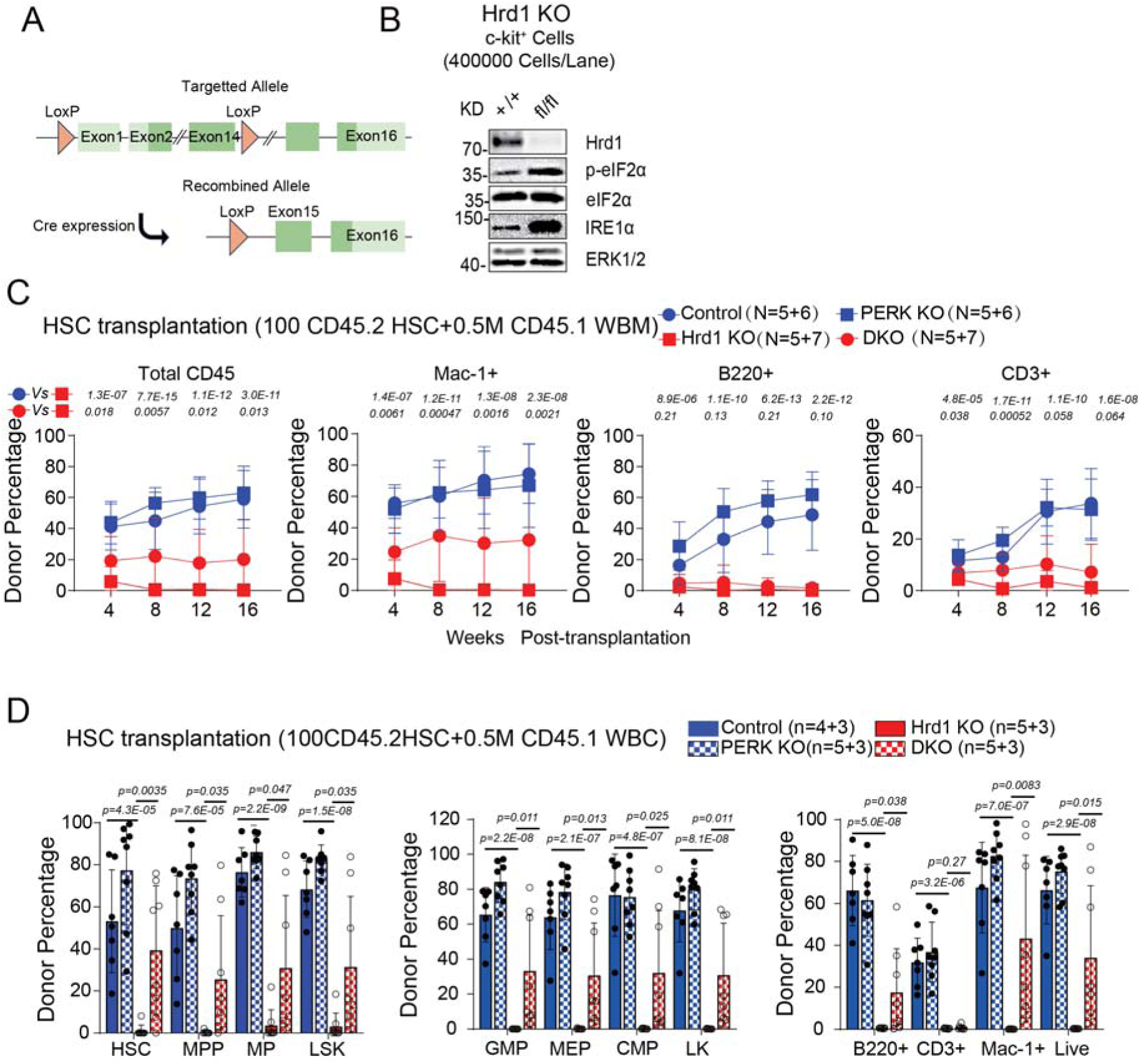
PERK knockout rescued HSC defects after ER stress induced by Hrd1 knockout. **A)**. Schematic representation of Floxed allele and wild type allele of *Hrd1 in* the conditional Hrd1 knockout mice. **B**). Western blot of Hrd1, IRE1α, total and phosphor-eIF2α in HSPCs of *Vav-cre^+^; Hrd1 ^fl/fl^* (Hrd1 KO) and control (*+*/*+*) mice. ERK level was used as loading control. Competitive repopulation assay with 100 FACS*-*purified HSCs from CD45.2 *Vav-cre^-^; Hrd1 ^fl/fl^*(control), *Vav-cre^+^; PERK ^fl/fl^*(PERK KO), *Vav-cre^+^; Hrd1 ^fl/fl^*(Hrd1 KO) ; *Vav-cre^+^; PERK ^fl/fl^; Hrd1 ^fl/fl^* (DKO) mice transplanted into irradiated receipt CD45.1 mice along with 0.5 million CD45.1 whole bone marrow competitors. The contribution of CD45.2 cells in total CD45^+^, myeloid (Mac1+), B (B220) and T (CD3+) cells from peripheral blood **(C)**, and in HSCs and other hematopoietic populations in the bone marrow (**D**) was analyzed in transplant recipients. Data represent mean±s.d. Each replicate represents a single mouse from two independent experiments.

Hrd1 KO mice displayed a similar phenotype as the Sel1L KO mice in reduction of bone marrow cellularity, spleen size and cellularity and thymus size and cellularity (**Supplementary Figure 4A-E**). The frequency and number of HSCs, MPPs, and LSKs are similarly decreased in both bone marrow (**Supplementary Figure 4F-G**) and the spleen (**Supplementary Figure 4H-I**). We also observed a similar myeloid biased lineage differentiation in the bone marrow and spleen (**Supplementary Figure 4J-M**). More importantly, there was a significantly increased phosphorylation of eIF2α, indicating activation of PERK signaling (**Figure 4B**) and accumulation of IRE1α, a substrate of ERAD^31^ (**Figure 4B**) in Hrd1 KO cells. Taken together, our results suggest that ERAD deficiency, via loss of either Sel1L or Hrd1, induces activation of ER stress and activates UPR PERK signaling.

### PERK is required for HSC dysregulation under ER stress induced by ERAD deficiency

Given that ERAD deficiency leads to the activation of all three branches of the UPR, we aimed to identify if a specific branch contributes to the loss of HSCs under these conditions. We generated *Mx1-cre^+^*; *Sel1L ^fl/fl^*; *PERK ^fl/fl^*(Sel1L/PERK double KO), *Mx1-cre^+^*; *Sel1L ^fl/fl^*; *IRE1*α *^fl/fl^* (Sel1L/IRE1α double KO), and *Mx1-cre^+^*; *Sel1L ^fl/fl^*; *ATF6 ^fl/fl^* (Sel1L/ATF6 double KO) mice. Knockout of PERK but not ATF6 or IRE1α in Sel1L KO mice resulted in a complete rescue of the reduction in HSC frequency (**Supplementary Figure 3J-K** and **Supplementary Figure 5F**) and number observed in Sel1L KO alone (**Supplementary Figure 5E**), suggesting that the PERK branch of the UPR contributes to Sel1L KO-mediated HSC depletion. We confirmed that PERK was deleted and phosphorylation of eIF2α decreased in Sel1L/PERK double KO mice (**Supplementary Figure 6A-B**). These results again suggested a PERK- dependent phosphorylation of eIF2α leads to HSC depletion in Sel1L knockout HSCs.

To determine if PERK also contributes to the impaired function of ERAD-deficient HSCs, we transplanted 5x10^5^ CD45.2*^+^* whole bone marrow cells from *Mx1-cre^+^*; *Sel1L ^fl/fl^*; *PERK ^fl/fl^*, *Mx1-cre^+^*; *Sel1L ^fl/fl^, Mx1-cre^+^; PERK ^fl/fl^*, or control (*Mx1-cre ^-^*; *Sel1L ^fl/fl^*; *PERK ^fl/fl^*) mice (without pIpC injection), along with 5x10^5^ CD45.1*^+^* competitor cells, into lethally irradiated CD45.1*^+^* recipients as previously described ^13^. Consistent with our previous report, Sel1L deletion induced by pIpC injection resulted in a significant depletion of Sel1L KO donor chimerism in the peripheral blood (**Figure 3C-F**). PERK KO was able to rescue peripheral blood chimerism (**Figure 3C-F**) up to 16 weeks.

More importantly, bone marrow chimerism analysis showed PERK knockout resulted in a complete rescue of chimerism loss in HSCs, MPPs, progenitors, myeloid cells but not B cell in Sel1L knockout (**Figure 3G**), indicating a cell type-specific regulation by PERK. These results suggest that PERK signaling contributes to the loss of HSC reconstitution potential induced by ERAD dysfunction.

As Hrd1 KO mice phenocopy the defects observed in Sel1L KO mice. To determine the effects of PERK KO on HSC dysregulation induced by Hrd1 loss, we generated *Vav-Cre^+^*; *Hrd1 ^fl/fl^; PERK ^fl/fl^, Vav-Cre^+^; Hrd1 ^fl/fl^*, *Vav-Cre^+^*; *PERK ^fl/fl^*, or control (*Vav-Cre ^-^*; *Hrd1 ^fl/fl^*; *PERK ^fl/fl^*) mice. Again, PERK KO decreases the phosphorylation of eIF2α in Hrd1 KO mice (**Supplementary Figure 6C**). Furthermore, similar to the results in PERK/Sel1L double knockout mice, the depletion of HSCs induced by Hrd1 KO was rescued by PERK deletion at the steady stage (**Supplementary Figure 6D-E**), confirming the significant contribution of PERK to HSC depletion upon ERAD deficiency at steady state.

To further assess the contribution of PERK to ERAD deficiency-mediated HSC impairment, competive transplantations were done with FACS-purified HSCs from *Vav-Cre^+^; Hrd1 ^fl/fl^; PERK ^fl/fl^*(Hrd1/PERK DKO), *Vav-Cre^+^*; *Hrd1 ^fl/fl^* (Hrd1 KO), *Vav-Cre^+^*; PERK *^fl/fl^*(PERK KO), or control (*Vav-Cre ^-^*; *Hrd1 ^fl/fl^*; *PERK ^fl/fl^*). Again, Hrd1 KO HSCs failed to provide any long-term reconstitution in transplant recipients (**Figure 4C**). PERK KO, on the other hand, significantly rescues the defect of HSC reconstitution (**Figure 4C**). Consistently, the levels of donor HSCs in the bone marrow of Hrd1 KO transplant recipients were significantly improved by PERK KO (**Figure 4D**). Thus, PERK signaling is activated in Hrd1 KO HSCs and is required for HSC depletion and defects induced by Hrd1 KO.

### PERK-mediated rescue of HSC defects in Hrd1 knockout depends on its kinase activity

PERK can function through both kinase-dependent and kinase-independent mechanisms to modulate cell fate in response to ER stress^19,32,33^. To assess whether PERK-mediated HSC dysregulation under ER stress induced by ERAD deficiency is dependent on kinase activity, we generated an in *situ* inducible knock-in of a kinase dead mutant^32^ strain, PERK K618A*^fl/fl^* (See methods for description, **Figure 5A**). As expected, both PERK knockout and PERK K618A knockin completely blocked the phosphorylation of eIF2α. Most importantly, unlike PERK knockout, the PERK kinase-dead mutation did not alter the protein level of PERK (**Figure 5B**).

**Figure 5.**
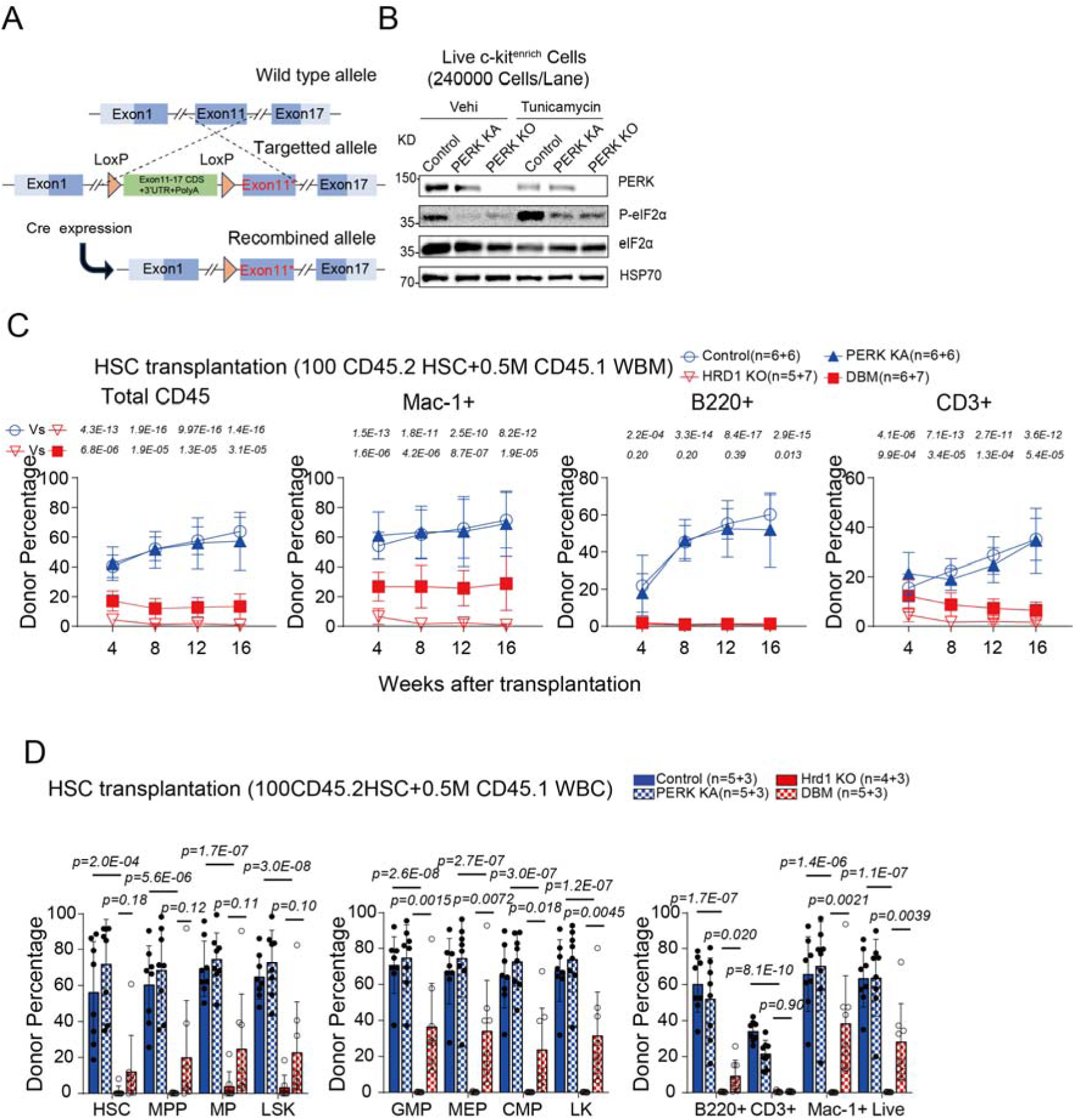
The effects of PERK on stressed HSCs depends on its kinase activity. **A).** Schematic representation of the knock-in allele of *PERK K618A***. B**). Western blot of p-eIF2α in live c-kit^+^ cell purified from *Vav1-cre^+^; PERK K618A^fl/fl^* (PERK KA) and *Vav1-cre^+^; PERK^fl/fl^* (PERK KO) mice, treated with DMSO (vehi) or tunicamycin for 4 hours. **C-D**). Competitive repopulation assay with 100 FACS*-*purified HSCs from CD45.2 *Vav-cre^+^; Hrd1 ^fl/fl^* (Hrd1 KO), *Vav-cre^+^; PERK K618A^fl/fl^* (PERK KA), *Vav-cre^+^; Hrd1 ^fl/fl^* (fl/fl) ; *PERK K618A^fl/fl^* (DBM) or control (*+*/*+*) mice transplanted into irradiated receipt CD45.1 mice along with 0.5 million CD45.1 whole bone marrow competitors. The contribution of CD45.2 cells in total CD45^+^, myeloid (Mac1^+^), B (B220^+^) and T (CD3^+^) cells from peripheral blood **(C)**, and in HSCs and other hematopoietic populations in the bone marrow (**D**) was analyzed in transplant recipients. Data represent mean±s.d. Each replicate represents a single mouse from two independent experiments.

We then evaluated the role of the kinase-dependent function of PERK in HSCs and hematopoiesis by crossing the *Mx1-cre^+^*mice with *PERK K618A ^fl/fl^* mice. PERK K618A KI phenocopy PERK KO mice in body weight, bone marrow, and spleen cellularity, as well as similar spleen and thymus weights and cellularity (**Supplementary Figure 7A-F)**, as well as all the hematopoitic population distribution(**Supplementary Figure 7G-R**). This was further confirmed by Vav1-cre-driven PERK K618A KI (**Supplementary Figure 8A-R**).

Next, we determined the impact of PERK kinase activity on HSC dysregulation induced by Hrd1 loss. We generated *Vav-Cre^+^*; *Hrd1 ^fl/fl^; PERK K618A ^fl/fl^*, *Vav-Cre^+^*; *Hrd1 ^fl/fl^, Vav-Cre^+^; PERK K618A ^fl/fl^*, and control mice. The depletion of HSCs, but not MPPs, induced by Hrd1 KO was rescued by PERK K618A KI at a steady state (**Supplementary Figure 9E-F**), underscoring the significant role of PERK kinase activity in HSC depletion upon ERAD loss. To further evaluate the function of individual HSCs, we did competitive transplantation with 100 FACS-purified HSCs from *Vav-Cre^+^; Hrd1 ^fl/fl^; PERK K618A ^fl/fl^, Vav-Cre^+^*; *Hrd1 ^fl/fl^, Vav-Cre^+^; PERK K618A ^fl/fl^,* and control mice. PERK K618A KI improves HSCs reconstitution ability observed in Hrd1 KO HSCs (**Figure 5C**). However, the levels of donor HSCs, progenitors, and lineage cells in the bone marrow of Hrd1 KO transplant recipients were not significantly rescued by PERK K618A KI (**Figure 5D**). Thus, PERK kinase activity plays a critical role in depleting Sel1L-Hrd1 ERAD KO HSCs in steady-state hematopoiesis.

### PERK depletes damaged HSCs without exacerbating ER stress

In the context of canonical apoptotic ER stress, PERK-eIF2α exacerbates ER stress by promoting protein synthesis and folding, eventually leading to ROS accumulation and cell death^17^. Although we did not observe any increase in ROS or cell death in Sel1L KO HSCs, it is still possible that PERK KO may rescue the defects of ERAD-deficient HSCs by reducing ER stress to a lower intensity. If this were true, we would expect lower ER stress intensity in Sel1L/PERK double KO. However, our results showed that both IRE1α-mediated Xbp1 splicing and downstream target Grp78^34^ levels were unchanged in PERK/Sel1L double knockout (**Supplementary Figure 10A-B**), suggesting that PERK-mediated rescue of the defects of Sel1L KO HSCs is not by reducing ER stress induced by Sel1L KO.

### PERK and ERAD deficiency collaborate to activate Rheb and promote mTOR signaling

Our previous studies suggest that ERAD deficiency leads to reduced HSC reconstitution by promoting HSC proliferation and differentiation^13^. To study the role of PERK in this process, we assessed the proliferation of HSCs in Sel1L/PERK double KO as well as Sel1L KO mice by Ki67 staining (**Figure 6A**). While PERK KO alone did not affect HSC proliferation, it completely blocked the increased proliferation of Sel1L KO HSCs. These results suggest that PERK depletes HSC via the promotion of HSC proliferation and loss of self-renewal.

**Figure 6.**
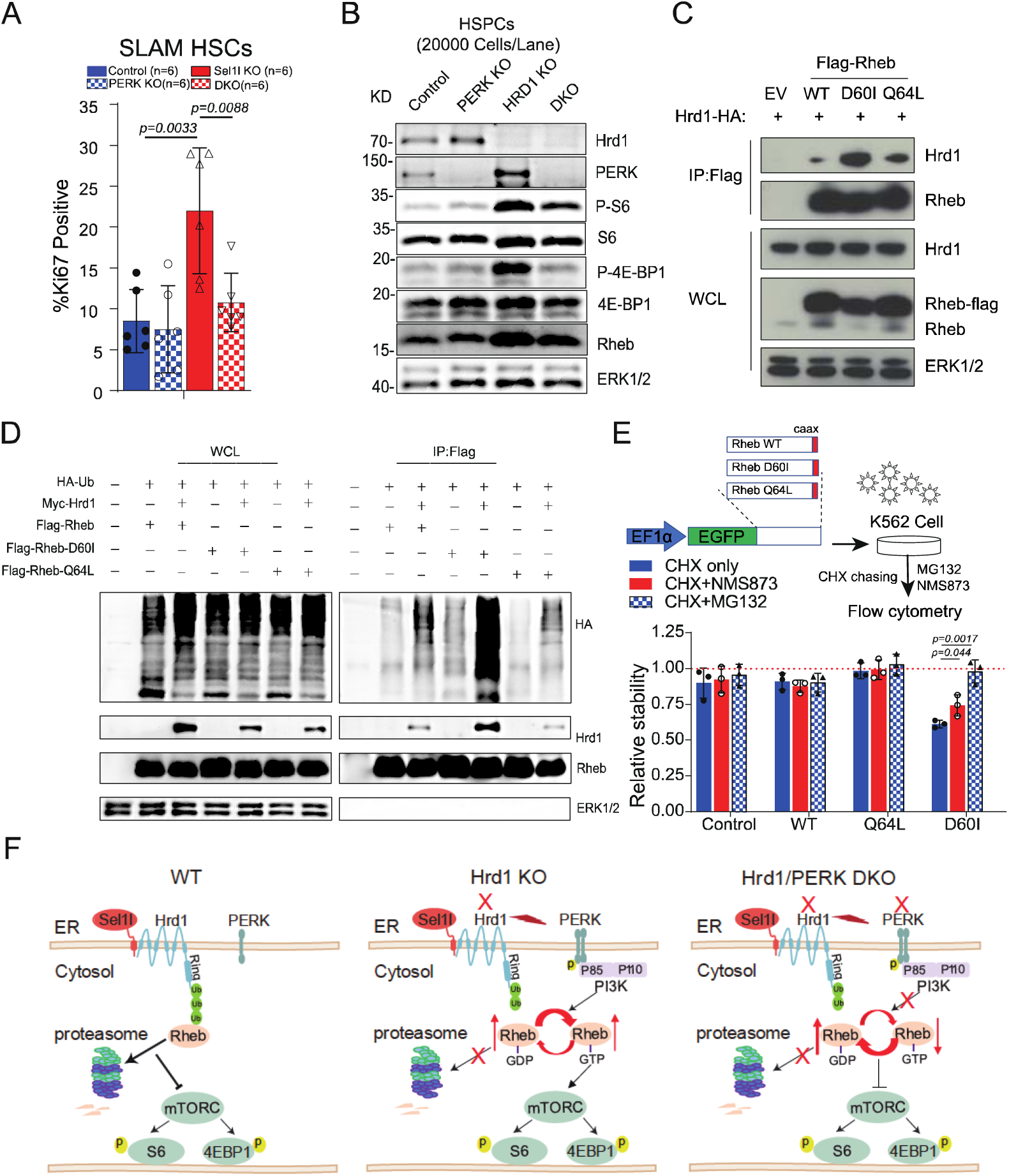
PERK and ERAD collaborate to activate mTOR to promote HSC proliferation. **A).** Ki67 staining of SLAM HSCs from 6-8 week old *Mx1-cre^-^; Sel1L ^fl/fl^* (WT), *Mx1-cre^+^; PERK ^fl/fl^* (PERK), *Mx1-cre^+^; Sel1L ^fl/fl^* (Sel1L) ; *Mx1-cre^+^; PERK ^fl/fl^; Sel1L ^fl/fl^* (DKO) mice 2 weeks after pIpC injection. **B**). Western blot of ERAD, UPR, mTORC1/2 proteins in HSPC (LSK) isolated from *Vav-cre^-^; Hrd1 ^fl/fl^* (control), *Vav-cre^+^; PERK ^fl/fl^* (PERK KO), *Vav-cre^+^; Hrd1 ^fl/fl^* (Hrd1 KO) ; *Vav-cre^+^; PERK ^fl/fl^; Hrd1 ^fl/fl^* (DKO) mice. Lysates from equal number of cells were loaded for each lane. **C-D)**. HEK 293T cells were transfected with constructs that overexpress HA-tagged ubiquitin, FLAG-tagged Rheb*-*WT; Rheb-D60I; Rheb-Q64L and Myc*-*tagged Hrd1. Rheb proteins were immunoprecipitated by FLAG antibody and Hrd1 binding to Rheb isoforms (**C)** and ubiquitinated Rheb (**D**) were detected by western blot. **E**). After GFP-fused Rheb and its mutants were stably overexpressed in the K562 cell line, the stability of each mutant was assessed by cycloheximide (CHX) treatment-protein stability tracking experiment. F). Schematic model of the collaboration of PERK and ERAD in regulating mTOR signaling under ER stress. Data represent mean±s.d. Each replicate represents a single mouse from an independent experiment. Two way Anova test was used for statistical analysis.

Our previous study suggested accumulation of Rheb contributes to mTOR activation and promotes hyper-proliferation of HSCs in Sel1L knockout HSCs^13^. Consistent with this, we observed that Hrd1 knockout led to significant mTORC1 signaling activation, as demonstrated by increased phosphor-S6 and phosphor-4EBP1 (**Figure 6B**).

Interestingly, although these increases were significantly blocked in Hrd1/PERK double KO (**Figure 6B**), Rheb protein was only slightly reduced in PERK/Hrd1 double KO as compared to Hrd1 KO (**Figure 6B**). This raises the possibility that the ERAD deficiency-mediated accumulation of Rheb alone was insufficient to induce mTORC1 signaling^35^.

Rheb is a small GTPase and cycles between an inactive Rheb-GDP bound conformation and an active Rheb-GTP bound conformation^36^. Upstream PI3K signaling leads to the activation of AKT, which promotes Rheb GTP-bound active conformation by inhibiting TSC complex^37^. We hypothesize that ERAD may preferentially degrade Rheb-GDP rather than Rheb-GTP, therefore ERAD insufficiency alone leads to accumulation of inactive Rheb-GDP and is not sufficient to activate Rheb. Since PERK has been shown to activate AKT in a p85-dependent manner upon ER stress^38^, the activation of PERK upon ERAD insufficiency may stimulate GDP-GTP conversion of the accumulated GDP-Rheb in Hrd1 KO, thereby enhancing the mTORC1 signaling. If this is true, we expect Hrd1 to preferentially bind GDP-bound Rheb. To test this, we transfected 293T cells with FLAG-tagged Rheb mutants that form GDP-bound (D60I) ^39^ or GTP-bound form Rheb (Q64L) ^40^, together with HA-tagged Hrd1, and performed co-immunoprecipitation assay with FLAG antibody to monitor levels of coIPed Hrd1 protein with these Rheb mutants. This showed that despite the lower expression of Rheb-GDP (Rheb-D60I) compared to both Rheb-GTP (Rheb-Q64L) and wild-type, the Rheb-GDP (Rheb-D60I) showed a much higher binding preference for Hrd1 compared to wild-type or Rheb-GTP (Q64L) (**Figure 6C**).

We next evaluated whether Hrd1 expression leads to differential ubiquitination of Rheb isoforms D60I and Q64L. We transfected 293T cells with FLAG-tagged Rheb mutants, together with HA-tagged ubiquitin and Myc-tagged Hrd1, and levels of immunoprecipitated Rheb proteins with FLAG antibody were monitored. As expected, we observed higher ubiquitination of Rheb-GDP^41^ (Rheb-D60I) by Hrd1 compared to other Rheb-GTP (Rheb-Q64L) and wild-type Rheb (**Figure 6D**), suggesting ERAD preferentially degrade GDP-bound Rheb.

To further confirm that this is tightly linked to the degradation of Rheb isoforms, we analyzed the stability of Rheb isoforms (**Figure 6E**). After protein synthesis was blocked by Cycloheximide (CHX), wild-type and GTP-bound Rheb were stable and did not show a change in their protein levels. However, Rheb-GDP (Rheb-D60I) showed a 50% reduction in its protein level suggesting that inactive GDP-bound Rheb is rapidly degraded (**Figure 6E**). This reduction in stability could be blocked by inhibition of ERAD downstream either via VCP (NMS-873^42^) or proteasome (MG-132^43^) inhibitors (**Figure 6E**). Collectively, these results suggest a stepwise activation of mTORC1 upon ERAD loss in HSCs (**Figure 6F**): first, the ERAD substrate Rheb-GDP accumulates due to the loss of Hrd1. Second, the accumulation of misfolded proteins activates PERK, which activates AKT via a P85-dependent manner and promotes Rheb-GDP to Rheb-GTP transition to fully activate mTORC1 signaling.

## Discussion

PERK has been extensively studied in ER stress^19^, and a previous study suggested that after ER stress-induced *in vitro* by tunicamycin or thapsigargin, PERK is activated to induce apoptosis to eliminate damaged HSCs^20^. However, tunicamycin or thapsigargin treatment induces severe ER stress, which may not reflect physiological conditions *in vivo*. The role of PERK *in vivo* in HSC has not been investigated. In this study, we report PERK is dispensable for hematopoiesis under steady-state conditions *in vivo*. However, upon ER stress induced by ERAD deficiency via Sel1L or Hrd1 knockout, PERK is activated to promotes HSC depletion. We previously reported that the levels of ERAD-related genes and ERAD activity are highest in dormant and quiescent HSCs and significantly decrease when HSCs exit quiescence and undergo proliferation and differentiation. Decrease of ERAD activity, therefore, occurs naturally during the transition from HSC quiescence to proliferation and activation, which represents a physiological condition with increased ER stress *in vivo*.

The canonical PERK branch of UPR is initiated by PERK phosphorylation of eIF2α, which leads to the inhibition of global protein translation and promotion of stress translation of a subset of proteins including ATF4^21^. ATF4 further activates transcription of CHOP that promotes the expression of *Dr5*^28^ *and Puma*^29^, both of which are sufficient to induce apoptosis^28,29^. Our previous report and studies here show that ERAD deficiency upon Sel1L or Hrd1 knockout did not induce apoptosis in HSCs *in vivo* but instead induced hyper-proliferation and differentiation leading to HSC depletion. PERK signaling therefore plays important role in non-apoptosis ER stress and can eliminate damaged HSCs independently of apoptosis by promoting proliferation and differentiation.

Accumulating studies suggest that the functions of PERK are mediated by both kinase-dependent and independent mechanisms ^44,45^ ^33^. For example, PERK is involved in the formation and maintenance of ER-mitochondria contact sites independent of its’ kinease activity^32^. In this study, we report that the function of PERK in HSC depletion is dependent on PERK’s kinase activity, and a “kinase dead” PERK mutant rescued HSC defects in Sel1L or Hrd1 KO similarly to PERK knockout mutant. We did, however, observe differential rescues in different hematopoietic populations. For instance, the depletion of HSCs, but not MPPs, induced by Hrd1 KO was rescued by PERK K618A KI at steady state (**Supplementary Figure 9E-F**), suggesting a differential dependency on PERK kinase activity in HSCs and MPPs.

Mechanistically, the activation of mTORC1 and hyperproliferation of HSCs induced by ERAD deficiency require PERK. Knockout of PERK blocked mTORC1 activation and HSC hyperproliferation in Hrd1 KO HSCs. Interestingly, while PERK knockout blocked mTORC1 activation in Hrd1 KO HSCs, it only modestly reduced the levels of Rheb.

This suggests that an increased level of Rheb alone is not sufficient to activate mTORC1 and prompted us to investigate the interaction between ERAD and PERK in regulating Rheb and mTORC1 activity. Previous studies suggested that PERK activates mTORC1 via PI3K/Akt-mediated inhibition of TSC ^38^. We found that Hrd1 preferentially binds and degrades Rheb-GDP to Rheb-GTP and Rheb-GDP is much more unstable than Rheb-GTP. We propose the model that upon ERAD deficiency, the reduced degradation leads to the accumulation of Rheb-GDP, and ER stress induced PERK activation leads to the transformation of Rheb-GDP to Rheb-GTP via activated PI3K/Akt (**Figure 6F**). Intriguingly, a previous study reported that the

GTP-bound active Rheb interacts with PERK and promotes eIF2α phosphorylation in response to ER stress^46^. Thus, the crosstalks between the ER stress signaling and the PI3K/Akt/mTOR are more complicated and likely to be context-dependent.

In summary, we present comprehensive studies that illustrate the function of PERK signaling in HSCs *in vivo,* both under steady-state and stressed conditions. While PERK is dispensable for steady-state hematopoiesis, it is required to preserve HSC integrity by promoting HSC hyperproliferation and differentiation, thereby eliminating damaged HSCs under ER stress in a kinase-dependent manner. These effects are regulated by a collaboration of PERK with ERAD to modulate Rheb stability and activation to affect mTOR activity. Further studies are required to dissect and fully understand the crosstalks among these important pathways to regulate HSC functions.

## Supporting information

Supplementary Methods, Supplementary Figures,Supplementary FigureLegends

## Acknowledgments

This work was supported by the University of Michigan Protein Folding Disease Initiative. E.M.K. received funding from the Research Training Award for Fellows from the American Society of Hematology and the pre-Career Development Award from the National Oncology Program. Q.L. was supported by NIH/NHLBI (1R01HL174566 and 1R01HL150707); L.L. was supported by National Natural Science Foundation of China (No. 32100636), High-level Talent Research Startup Fund of Hunan University, Provincial Natural Science Foundation of Hunan (2023JJ30125), The Excellent Youth Foundation of Hunan Province (2024JJ2017),

## Author Contributions

M.Z., Q.P., E.K., performed most of the experiments, interpreted results and assisted with manuscript preparation. Z.S., S.L., and Z.Y. performed some of the experiments with help from L.L. X.W. contributed to the PERK K618A point mutation mouse generation. Y.M. contributed to the flow cytometry and isolation of hematopoietic cells.L.L and Q.L. conceived the project, designed experiments, interpreted results, and wrote the manuscript.

## Data availability

All other data supporting the findings of this study are available from the corresponding author upon reasonable request.

