## Supplementary Methods, Supplementary Figures,Supplementary FigureLegends for "PERK Signaling Maintains Hematopoietic Stem Cell Pool Integrity under Endoplasmic Reticulum Stress by Promoting Proliferation"

#### Mice

For reconstitution assays, adult C57BL/Ka-CD45.1: Thy-1.2 mice, aged at least 8 weeks at the time of irradiation, served as recipients. pIpC was prepared in PBS and administered via intraperitoneal injection at a dose of 0.5 μg/g body weight every other day for six doses, or 2.65 μg/g body weight for three doses. All mice were analyzed at 8-14 weeks of age, with a minimum of two weeks post-pIpC treatment, and were paired with age- and sex-matched controls. Analyses were performed on samples from age- and sex-matched mice, incorporating both male and female subjects, with no observed differences between genders. Efforts were made to ensure equal representation of male and female mice in the analyses.

Heterozygous mice with a conditional point mutation in the Eif2ak3 gene (K618A), designated as PERK K618A $^{fl/-}$ mice, were generated at Shanghai Model Organisms. The kinase dead mutant strain, PERK K618A$^{fl/fl}$, which abolishes the kinase activity of PERK. Using the principle of homologous recombination, we performed flox modification on the Eif2ak3 (encoding PERK) gene. The Exon11 sequence was replaced by Loxp-Exon11-Exon17CDS-3'UTR-PolyA signal-Loxp-Exon11*(lysine to alanine substitution at amino acid 618; K618A). When Cre is expressed, "Exon11-Exon17CDS-3'UTR-PolyA signal" segment is deleted, enabling the expression of Exon11*(K618A) and the downstream sequence of Exons 12-17, therefore the PERK K618A mutant is expressed. These mice were maintained on a C57BL/6 background.

#### Flow cytometry and isolation of hematopoietic cells.

Bone marrow cells were isolated from the long bones (tibias and femurs) of mice by flushing with FACS Buffer (Hank's buffered salt solution without calcium or magnesium, supplemented with 2% heat-inactivated calf serum). The collected cells were triturated and filtered through a nylon mesh to obtain a single-cell suspension. Hematopoietic populations were analyzed using the antibody cocktails listed in **Supplementary Table 3**, following previously described protocols.

For the isolation of hematopoietic stem cells (HSCs), whole bone marrow cells were incubated with the antibody cocktails indicated in **Supplementary Table 3**. Following washing, the cells were incubated with anti-APC-conjugated paramagnetic microbeads. The microbead-bound (c-kit+) cells were then enriched using MS columns (Miltenyi Biotec). Non-viable cells were excluded from sorting and analyses using the viability dye 4′,6-diamidino-2-phenylindole (DAPI) at a concentration of 1 μg/ml. Cell analysis was performed using a BD Fortessa flow cytometer, while sorting was conducted with a BD Aria III flow cytometer. Additional information regarding the antibodies used can be found in **Supplementary Table 1**. Additional information of methods and materials can be found in **Supplementary methods** and **Supplementary Table 1-3**.

**Genotyping**

A total of 150 μL of 50 mM NaOH was added to a 200 μL tube containing either a 1-2 mm portion of mouse tail or at least 1 million cells from each mouse. The samples were then incubated at 90 °C for 1 hour to facilitate lysis. Following the heating step, 50 μL of Tris-HCl (pH 8.0) was added to each tube to neutralize the reaction. The samples were vortexed thoroughly to ensure complete mixing. Subsequently, 2 μL from each sample was utilized for the genotyping PCR reaction. Genotyping PCR was conducted using GoTaq® Green Master Mix according to the manufacturer's instructions. All primers used are listed in **Supplementary Table 2**.

**Competitive repopulation assay**

Adult recipient mice (CD45.1) were irradiated using Radiosource (RS2000 Pro) delivering approximately 75 rad $min^{-1}$ in a single dose of 750 rad. Following irradiation, cells were injected into the tail veins of anesthetized recipients. Blood samples were collected from the tail veins of recipient mice starting at 4 or 6 weeks post-transplantation and continuing for at least 16 weeks. The samples were treated with ammonium-chloride potassium red blood cell lysis buffer (8 g/L NH4Ac, 1 g/L KHCO3, 0.04 g/L EDTA) and stained with directly conjugated antibodies to CD45.2 (104), CD45.1 (A20), B220 (6B2), Mac-1 (M1/70), CD3 (KT31.1), and Gr-1 (8C5) to assess engraftment (refer to the Lineage Bleeding Cocktail in **Supplementary Table 3**). Chimerism of stem cells, progenitors, and lineage cells was analyzed at least 16 weeks after transplantation, unless otherwise specified. Additional information regarding the antibodies is provided in **Supplementary Table 1**.

**Immunoprecipitation**

Co-immunoprecipitation assays were performed in 293T cells. Briefly, the cells were lysed in lysis buffer (150 mM NaCl, 1 mM EDTA, 50 mM Tris-HCl pH 7.5 or 8.0, supplemented with protease and phosphatase inhibitors, 10 mM N-ethylmaleimide, and 1% Nonidet P-40 (NP-40)). Protein lysates were incubated with anti-Flag magnetic beads (Thermo Fisher A36798) or control beads. Immunocomplexes were precipitated using a Magnetic Separation Rack, washed, and then eluted by boiling for 10 minutes in 2× SDS sample buffer.

**Western blotting**

An equal number of LSK cells (20,000 cells) from each mouse were sorted directly into PBS and precipitated with trichloroacetic acid (TCA) to a final concentration of 10% TCA. Extracts were incubated on ice overnight and centrifuged for 10 minutes at 13000rpm at 4 °C. The supernatant was removed, and the precipitated pellets were washed twice with ice-cold acetone and airdried. The proteins in the pellets were solubilized with solubilization buffer (9 M urea, 2% Triton X-100, 1% DTT) before the addition of Omni-Easy™ Fast Protein Loading buffer. Proteins were separated on SuperPAGE™ 4-12% Bis-Tris Protein Gels and transferred to PVDF membranes. The antibodies used are listed in **Supplementary Table 1**.

**Protein aggregation Assay**

The protein aggregation assay was performed by collecting 20 million whole bone marrow cells and staining them with FACS antibodies against cell surface markers for HSCs and other hematopoietic populations, as described in **Supplementary Table 3**. After washing, the cells were resuspended and fixed with Cytofix/Cytoperm™ Buffer (BD Pharmingen™ BrdU Flow Kits) on ice for 10 minutes. The cells were then permeabilized using BD Cytoperm™ Permeabilization Buffer Plus (BD Pharmingen™ BrdU Flow Kits). Following a wash with 1X Perm/Wash™ Buffer (BD Pharmingen™ BrdU Flow Kits), the cells were resuspended in 1X Perm/Wash™ Buffer containing 1/5000 PROTEOSTAT® detection reagent and stained for 30 minutes. After another wash with FACS Buffer, the cells were resuspended in FACS Buffer containing 2.5 μg/ml DAPI. Aggregation was detected using the 582/15 (488) channel on a BD Fortessa flow cytometer.

**In vitro Annexin V staining of HSCs**

For in vitro staining, a minimum of 1000 hematopoietic stem cells (HSCs) or CD48-LSK cells were sorted into SF-03 medium supplemented with 100 ng/ml thrombopoietin (TPO) and stem cell factor (SCF). Cells were allowed to recover overnight before being treated with tunicamycin or thapsigargin for 18 hours. Subsequently, the cells were stained with APC-Annexin V (1:40 dilution) and 2.5 μg/ml DAPI.

**Ki67 Assay.**

10 million whole bone marrow cells were collected and stained with a cocktail of FACS antibodies, as detailed in **Supplementary Table 3**. Following washing with FACS buffer, cells were fixed and permeabilized according to the BrdU assay kit protocol. Subsequently, the cells were stained with FITC-conjugated Ki67 (BD Biosciences, 556026), and Ki67 levels were assessed using flow cytometry.

**CFU assay**

For colony formation assays, 200 SLAM HSCs were sorted directly into MethoCult® GF M3434 medium (Stem Cell Technologies, 3434) supplemented with 100 μl SF-03 medium containing 100 ng/ml TPO. The cells were mixed thoroughly by vortexing and then plated onto culture dishes. After 7 days of incubation, colonies were counted.

**Lentiviral Production and Transduction of Suspension Cells assay**

293T cells were transfected with a lentiviral expression system, including the appropriate packaging plasmid (pMD2.G) and helper plasmid (psPAX2). Following transfection, the viral supernatant was harvested and filtered through a 0.45 μm filter to remove cell debris. K562 cells were cultured in an appropriate medium until they reached the logarithmic growth phase. The filtered viral supernatant was supplemented with polybrene (10 μg/mL) and mixed with the suspension cells. The mixture was centrifuged at 2500 rpm at 37°C for 90 minutes to facilitate viral entry. Following centrifugation, the cells were cultured under standard conditions (37°C, 5% CO2). Fresh medium was typically replaced after 8-10 hours, with continued culture for an additional 24-48 hours to ensure effective viral invasion. After the incubation period, the medium was replaced with fresh medium containing blasticidin to select for successfully transduced cells.

**Cycloheximide (CHX) assay**

K562 cells were cultured in RPMI-1640 medium supplemented with 10% fetal bovine serum (FBS) at 37°C in a 5% $CO_2$ incubator, maintaining a density of 1–5 × $10^5$ cells/ml. A stock solution of cycloheximide (CHX) was prepared at a concentration of 10 mg/ml in DMSO. CHX was then diluted in fresh RPMI medium to a final concentration of 10–100 μg/ml and added to the K562 cells, with a control group receiving no CHX treatment. Cells were incubated for varying time points (e.g., 0, 1, 2, 4, and 8 hours) to assess the effects of CHX on protein levels over time. Following the incubation period, the media were removed, and the cells were washed twice with cold PBS.

**ER tracker, ROS staining**

20 million whole bone marrow cells were collected and stained with FACS antibodies against cell surface markers for HSCs and other hematopoietic populations, as detailed in **Supplementary Table 3**.

**RT-qPCR**

HSCs were FACS-sorted directly into 500 μl TRIzol™ Reagent, and RNA extraction was performed according to the manufacturer's instructions. To enhance RNA precipitation, 12 μl of Linear Acrylamide was added. Extracted RNA was dissolved in DPEC water and reverse-transcribed to cDNA using the High-Capacity RNA-to-cDNA™ Kit following the provided protocol. The resulting cDNA was utilized for qPCR with Power SYBR™ Green PCR Master Mix. All qPCR primers are listed in **Supplementary Table 2**.

**Quantification and statistical analysis**

All quantitative data are presented as mean ± standard deviation, unless stated otherwise. For statistical analysis, two-tailed Student's t-tests were employed for comparisons between two groups, while ANOVA was used for comparisons involving more than two groups, using Prism 7 (GraphPad Software). No randomization or blinding was applied in any experiments, and no experimental mice were excluded from the analysis. Sample sizes were determined based on experimental variation within control groups and are specified in the figure legends.

### Supplementary Figures

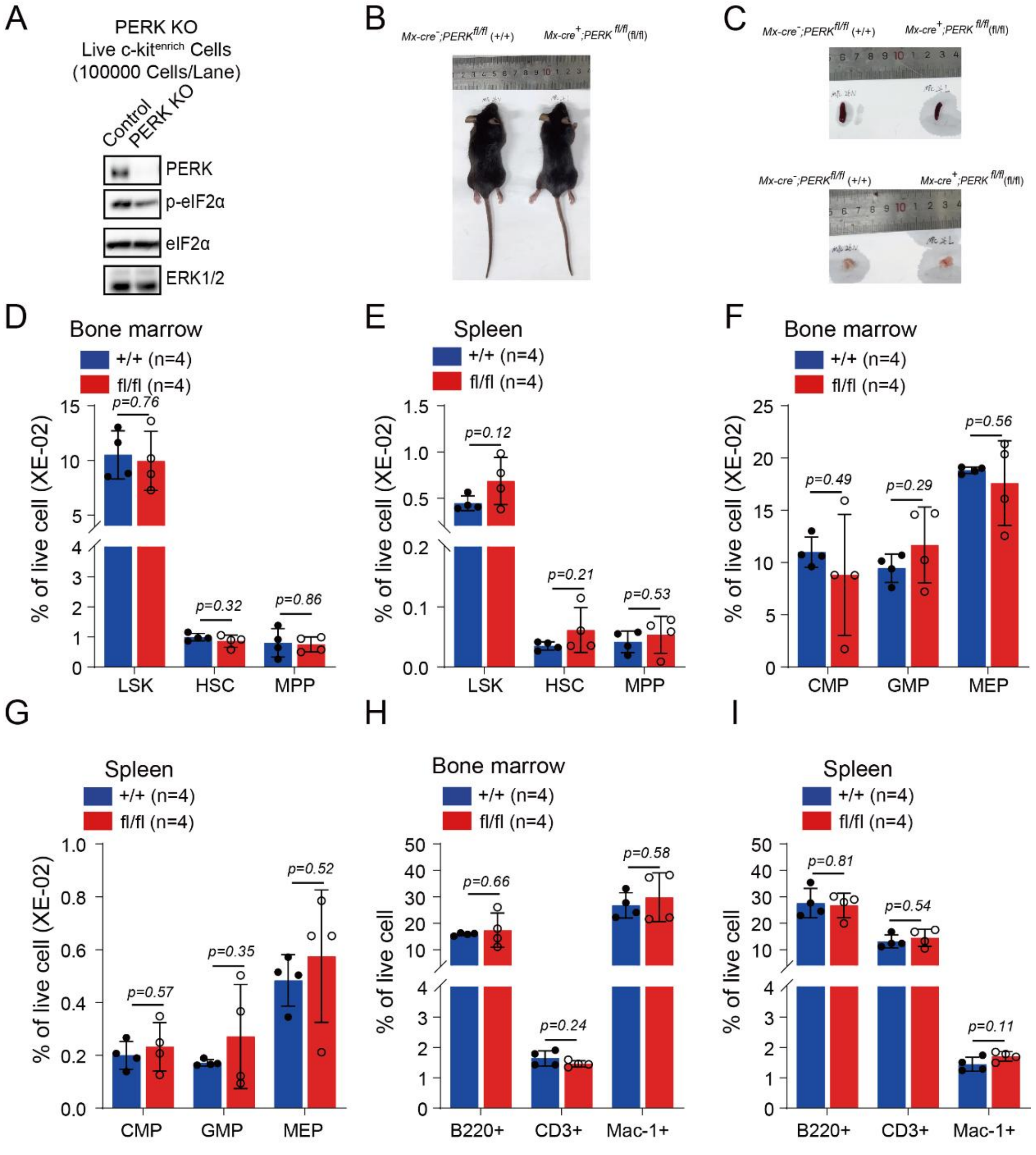


**Supplementary Figure 1. Steady state hematopoiesis in *Mx1-cre$^{+}$; PERK$^{fl/fl}$* mice. A)**. Western blot of PERK, phosphor-eIF2α, total eIF2α in live c-kit$^{+}$ cell**.** ERK was used as loading control. **B-C)**. Representative photos of the mice (**B**), spleen, and thymus **(C)** in mutants and age- and sex-matched controls (*Mx1-cre$^{-}$; PERK$^{fl/fl}$*). **D-I)**. Frequencies of hematopoietic populations in the bone marrow and spleen, 2 weeks after pIpC injection. Data represent mean±s.d. Each replicate represents a single mouse from an independent experiment. Two-sided student t-test was used for statistical analysis.

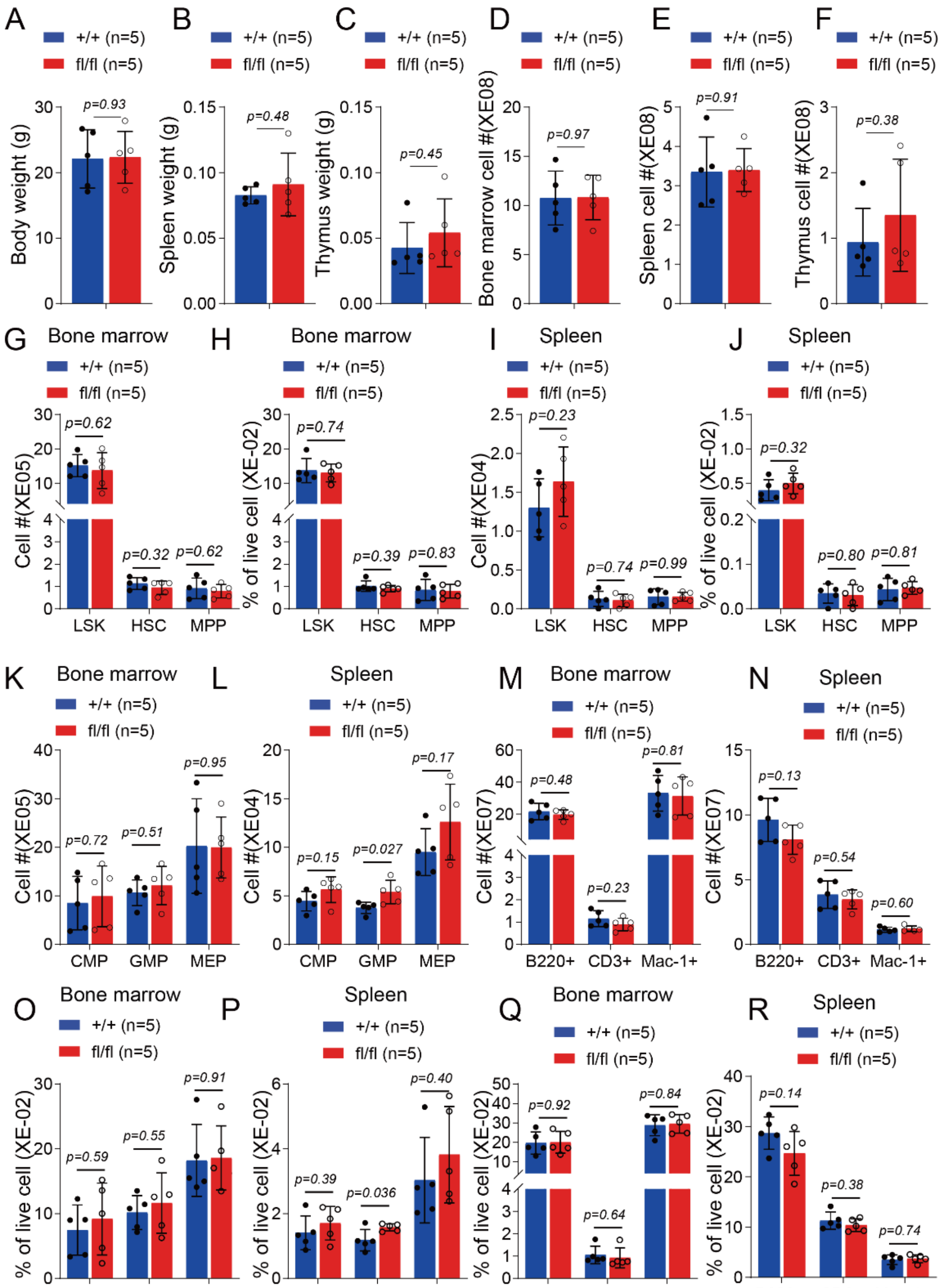


**Supplementary Figure 2. Steady state hematopoiesis in *Vav1-cre*$^{+}$*; PERK*$^{fl/fl}$ mice.**

6 to 8 weeks old *Vav1-cre*$^{+}$*; PERK*$^{fl/fl}$ (fl/fl) and *Vav1-cre*$^{-}$*; PERK*$^{fl/+}$ or *Vav1-cre*$^{-}$*; PERK*$^{fl/fl}$ (+/+) mice were analyzed for body weight (**A**), spleen (**B**) and thymus (**C**) weight, cellularity of bone marrow (**D**), spleen (**E**) and thymus (**F**), frequency and numbers of HSPC (**G-J**), numbers of lineage restricted progenitors (**K-L**) and mature blood cells (**M-N**), frequency of lineage

restricted progenitors (**O-P**) and mature blood cells (**Q-R**). Data represent mean±s.d. Each replicate represents a single mouse from an independent experiment. Two-sided student t-test was used for statistical analysis.

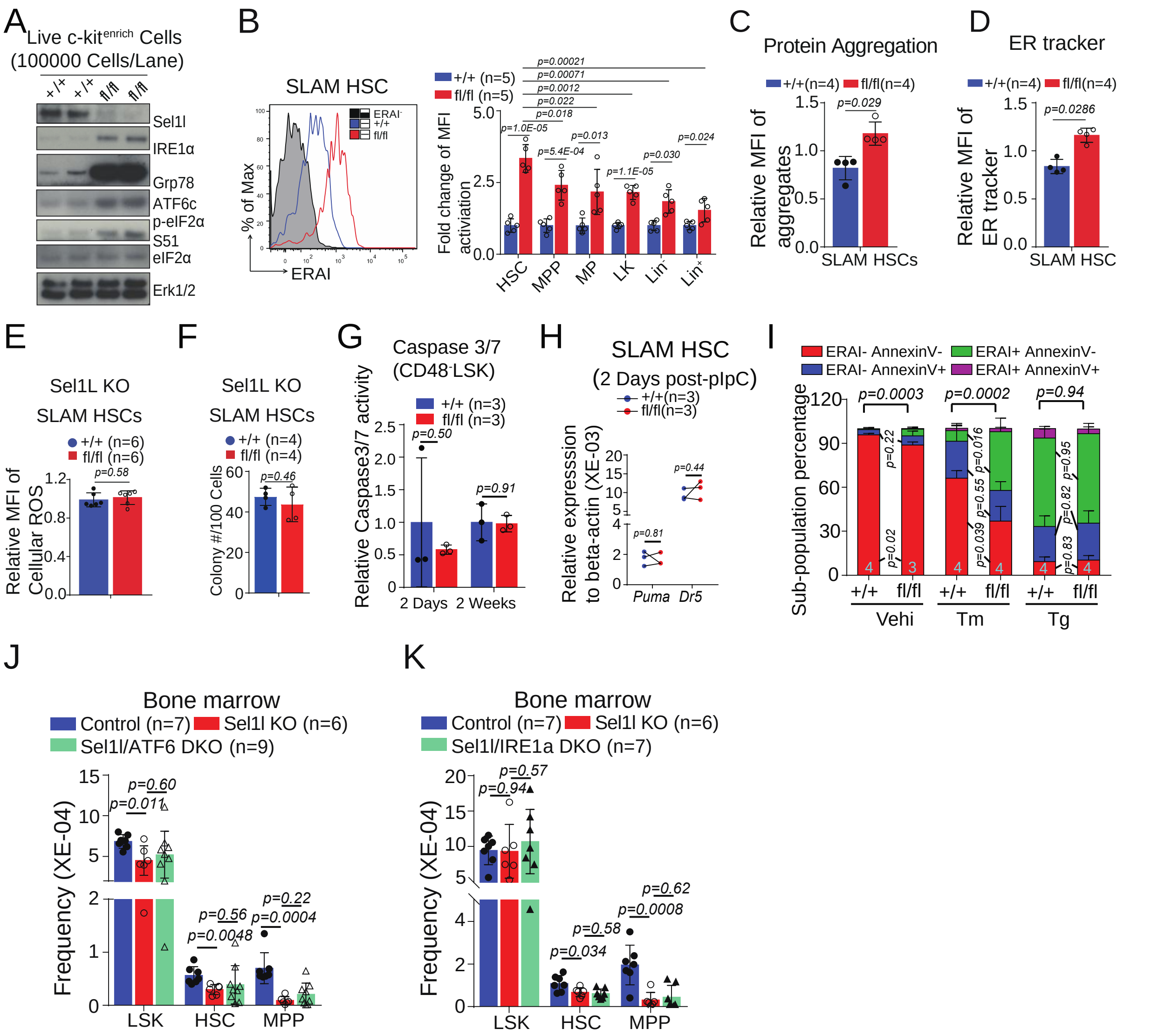


**Supplementary Figure 3. ERAD deficiency via Sel1L knockout induces non-apoptotic stress in HSCs. A).** Western blot of ER stress signaling targets in HSPCs from *Mx1-cre*$^{+}$*; Sel1L*$^{fl/fl}$ (fl/fl) or control (+/+) mice two weeks after pIpC injection. **B**). Two weeks after pIpC injection, ERAI in hematopoietic populations of *Mx1-cre*$^{+}$*; Sel1L*$^{fl/fl}$*; ERAI*$^{+}$ (fl/fl) and control (+/+) mice were detect by FACS. Data represent mean fluorescence intensity relative to *ERAI-* mice. **C-E**). Two weeks after pIpC injection, *Mx1-cre*$^{+}$*; Sel1L*$^{fl/fl}$ (fl/fl) or control (+/+) mice were analyzed for protein aggregates (by PROTESTAT, **C**) and ER volume (by ER-Tracker, **D**), and total ROS accumulation (**E**). **F**). Colony forming Units from 100 SLAM HSCs isolated from *Mx1-*

*cre*$^{+}$; *Sel1L*$^{fl/fl}$ (fl/fl) or control (+/+) mice. **G-I**). Levels of apoptosis measured by caspase 3/7 (**G**) or Annexin V staining (**I**) and apoptosis genes (by qPCR; **H**) in SLAM HSCs isolated from *Mx1-cre*$^{+}$; *Sel1L*$^{fl/fl}$ (fl/fl) or control (+/+) mice two weeks after pIpC injection. **J-K**). HSC frequencies in the bone marrow of Sel1L/ATF6 (**J**) or Sel1L/IRE1a (**K**) double knockout mice two weeks after pIpC injection**.** Data represent mean±s.d. Each replicate represents a single mouse from an independent experiment.

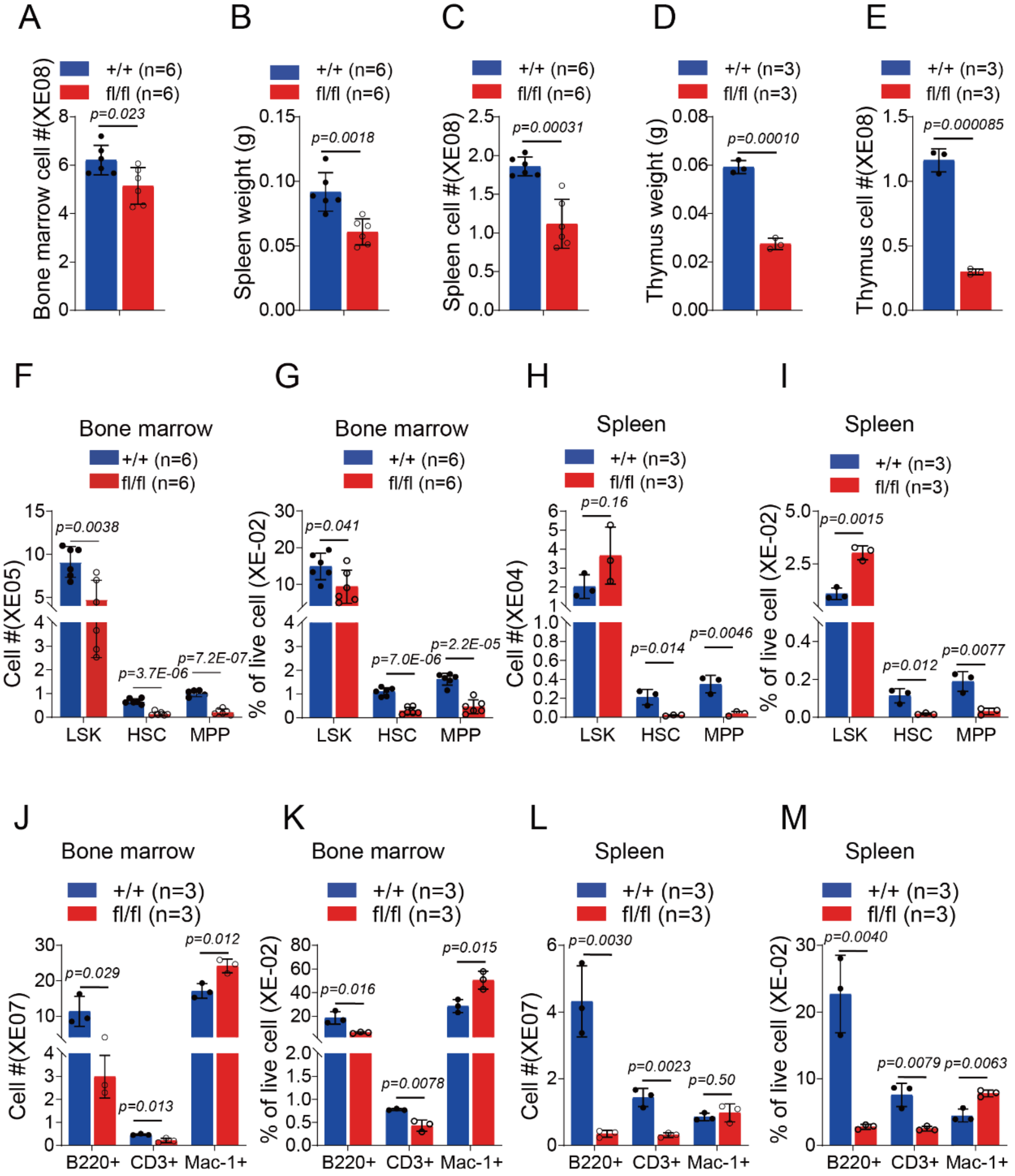


**Supplementary Figure 4. Steady state hematopoiesis in *Vav1-cre*$^{+}$; *Hrd1*$^{fl/fl}$ mice.**

6 to 8 weeks old *Vav1-cre*$^{+}$; *Hrd1*$^{fl/fl}$ (fl/fl) and *Vav1-cre*$^{-}$; *Hrd1*$^{fl/+}$ or *Vav1-cre*$^{-}$; *Hrd1*$^{fl/fl}$ (+/+) mice were analyzed for cellularity of bone marrow **(A)**, spleen weight **(B)** and cellularity **(C)**, thymus weight (**D**) and cellularity **(E)**, frequency and numbers of HSPC **(F-I)**, frequency and numbers of mature blood cells (**J-M**) in spleen and bone marrow. Data represent mean±s.d. Each replicate

represents a single mouse from an independent experiment. Two-sided student t-test was used for statistical analysis.

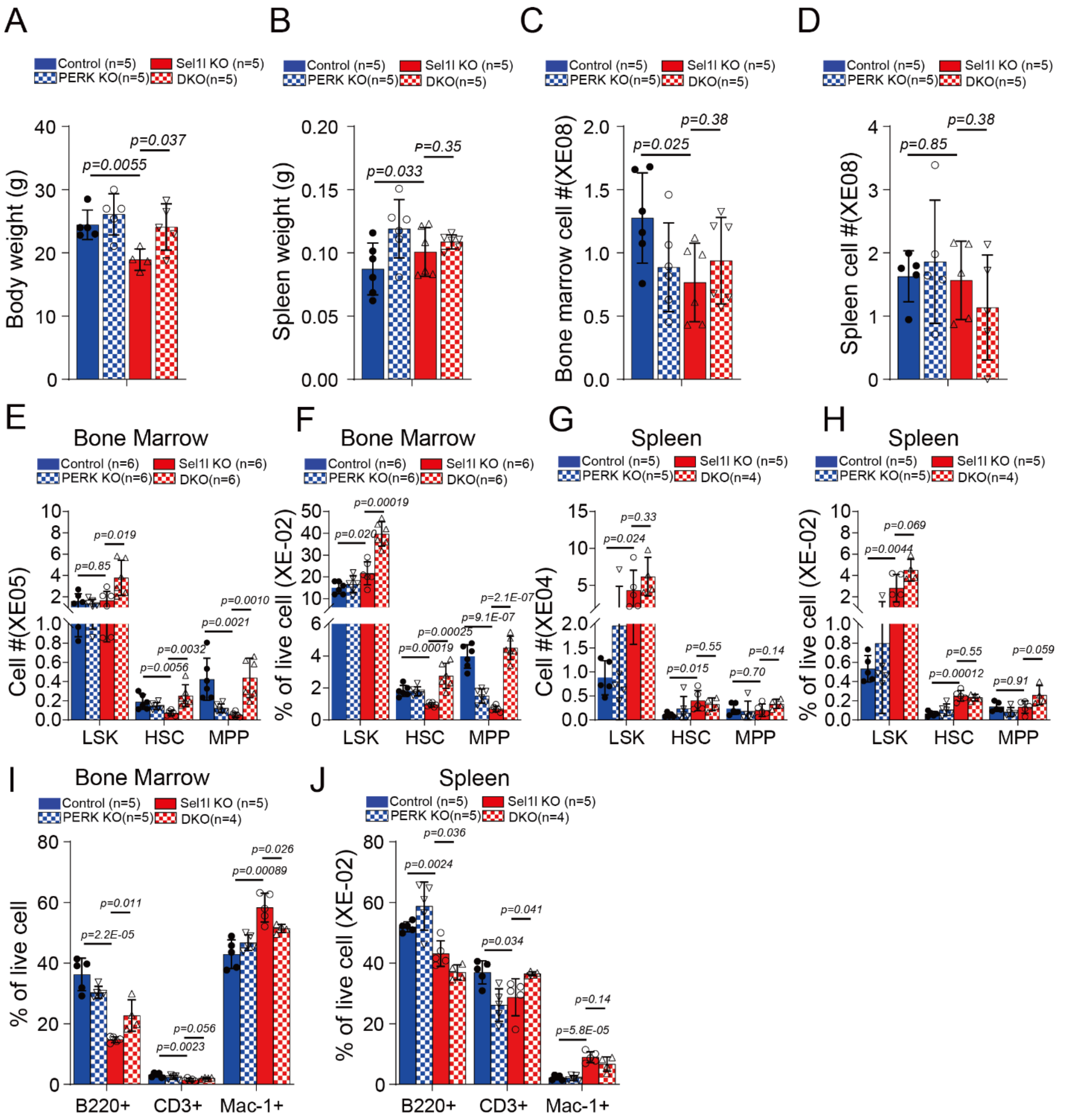


**Supplementary Figure 5. Steady state hematopoiesis in *Sel1L/PERK double knockout (KO)* mice.** *Mx1-cre*$^{+}$*; Sel1L*$^{fl/fl}$ (Sel1L KO) and *Mx1-cre*$^{+}$*; PERK*$^{fl/fl}$ (PERK KO)*; Mx1-cre*$^{+}$*; Sel1L*$^{fl/fl}$; *PERK*$^{fl/fl}$ (DKO), or control mice were analyzed for cellularity of body weight (**A**), spleen weight (**B**), bone marrow **(C)** and spleen cellularity **(D)**, frequency and numbers of HSPC in the bone marrow **(E-F)** and spleen **(G-H)**, frequency and numbers of mature blood cells in bone marrow

(**I)** and spleen (**J**). Data represent mean±s.d. Each replicate represents a single mouse from an independent experiment. Two-sided student t-test was used for statistical analysis.

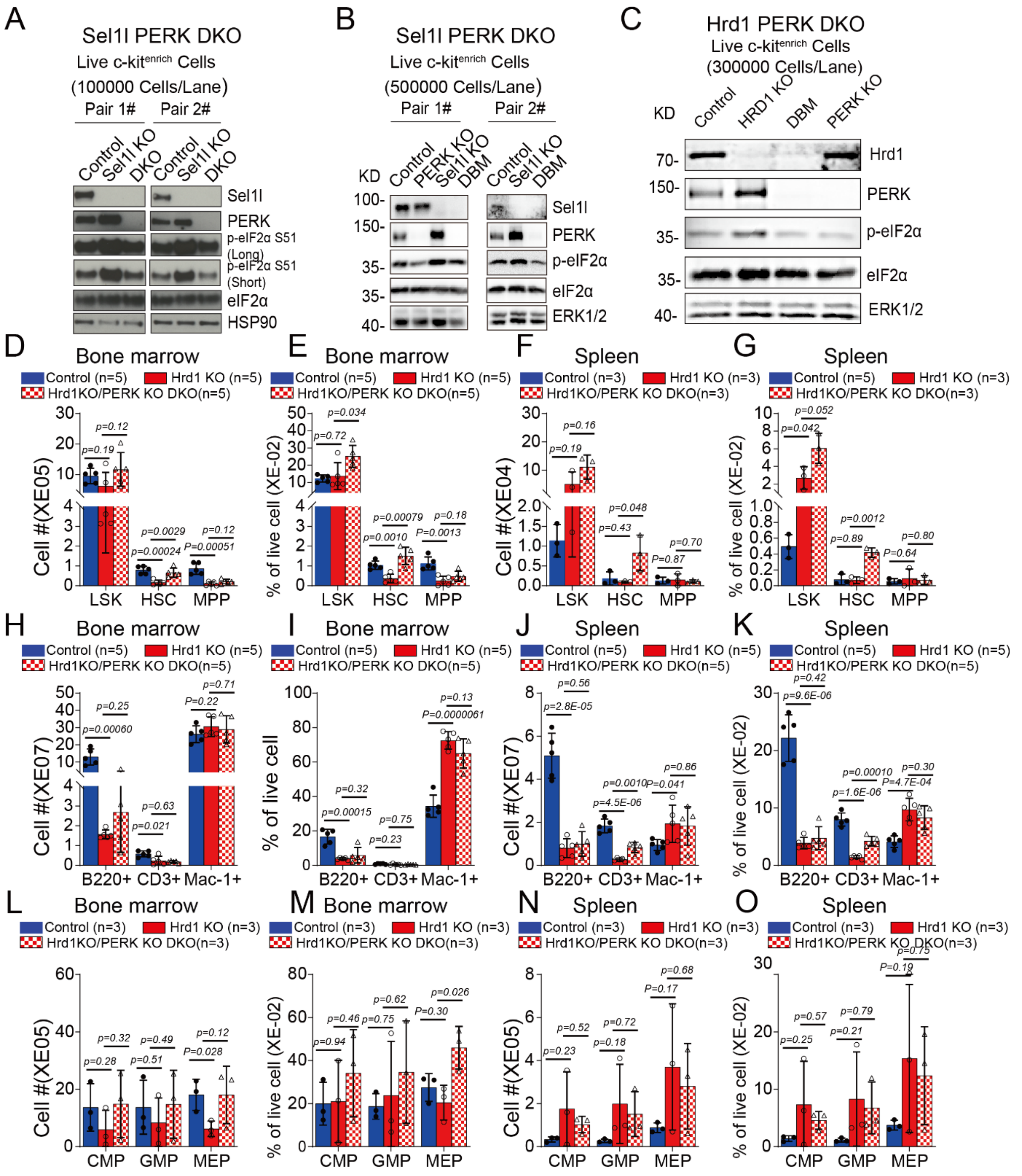


**Supplementary Figure 6. PERK rescue the depletion of Stressed HSCs**

Western blot of ERAD, PERK-eIF2α in *Mx1-cre*$^{+}$*; Sel1L*$^{fl/fl}$ (fl/fl)*; PERK*$^{fl/fl}$ (fl/fl) or control (+/+) mice live c-kit$^{+}$ cells **(A, B)**, and *Vav-cre*$^{+}$*; Hrd1*$^{fl/fl}$ (fl/fl); *PERK*$^{fl/fl}$ (fl/fl) or control (+/+) mice live c-kit$^{+}$ cells **(C)**. 6 to 8 weeks old *Vav-cre*$^{+}$*; Hrd1* $^{fl/fl}$ (fl/fl), *Vav-cre*$^{+}$*; Hrd1*$^{fl/fl}$ (fl/fl); *PERK*$^{fl/fl}$ (fl/fl) or control (+/+) mice were analyzed for frequency and numbers of HSPC **(D-G)**, mature blood cells

**(H-K)** and lineage restricted progenitors (**L-O**) in spleen and bone marrow. Data represent mean±s.d. Each replicate represents a single mouse from an independent experiment. Two-sided student t-test was used for statistical analysis unless specified.

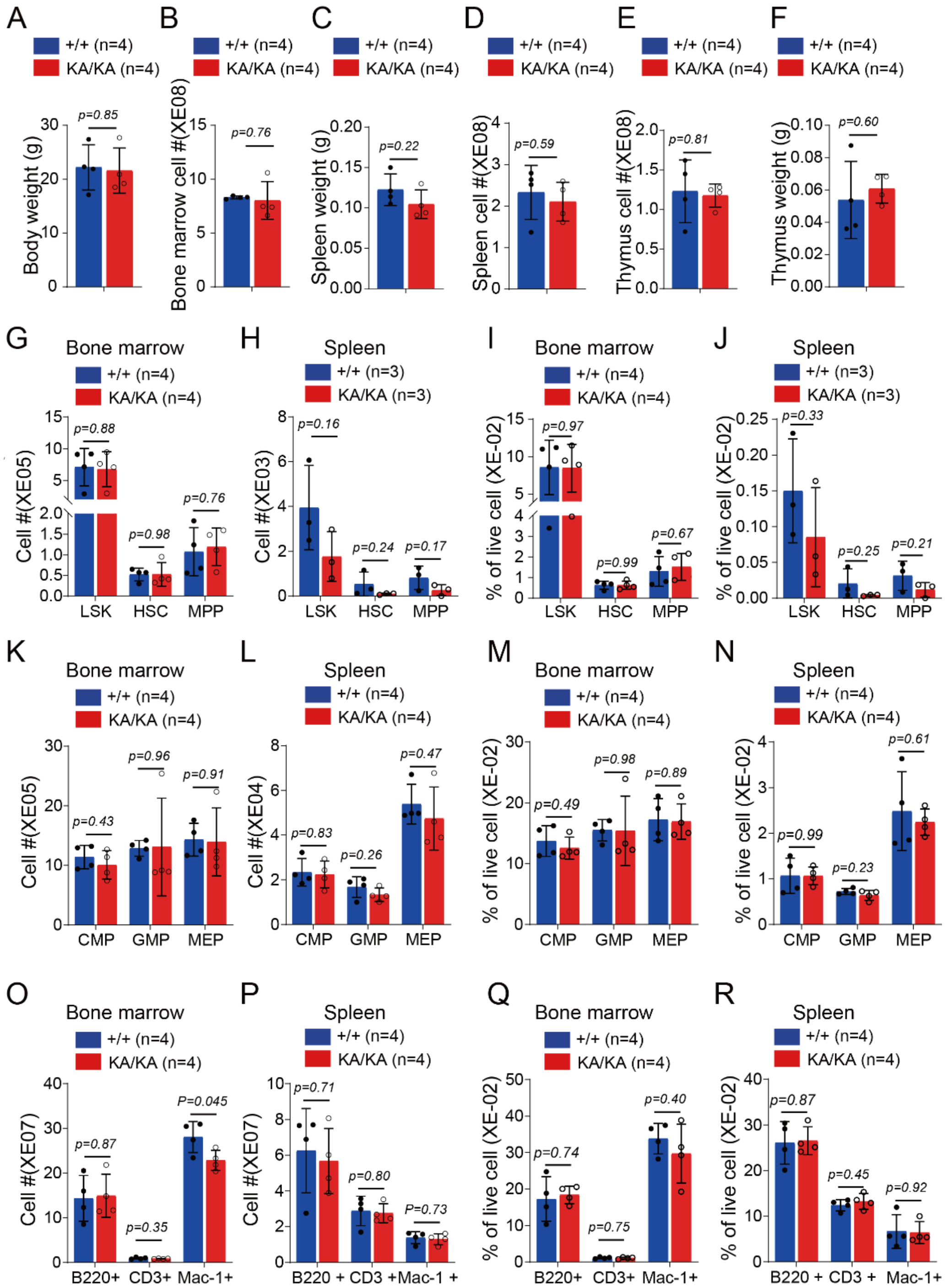


**Supplementary Figure 7. Steady state hematopoiesis in *Mx1-cre*$^{+}$*; PERK K618A*$^{fl/fl}$ mice**

6-8 weeks old *Mx1-cre*$^{-}$*; PERK K618A*$^{fl/fl}$ or and *Mx1-cre*$^{+}$*; PERK K618A*$^{fl/fl}$ (fl/fl) mice were injected with pIpC every other day for a total of 3 doses. Two weeks after pIpC injection, body weight **(A)**, spleen **(B)** and thymus **(C)** weight, cellularity of bone marrow **(D)**, spleen **(E)** and thymus **(F)**, frequency and numbers of HSPC **(G-J)**, lineage restricted progenitors **(K-N)** and mature blood cells **(O-R)** were analyzed. Data represent mean±s.d. Each replicate represents a single mouse from an independent experiment. Two-sided student t-test was used for statistical analysis unless specified. Data represent mean±s.d.

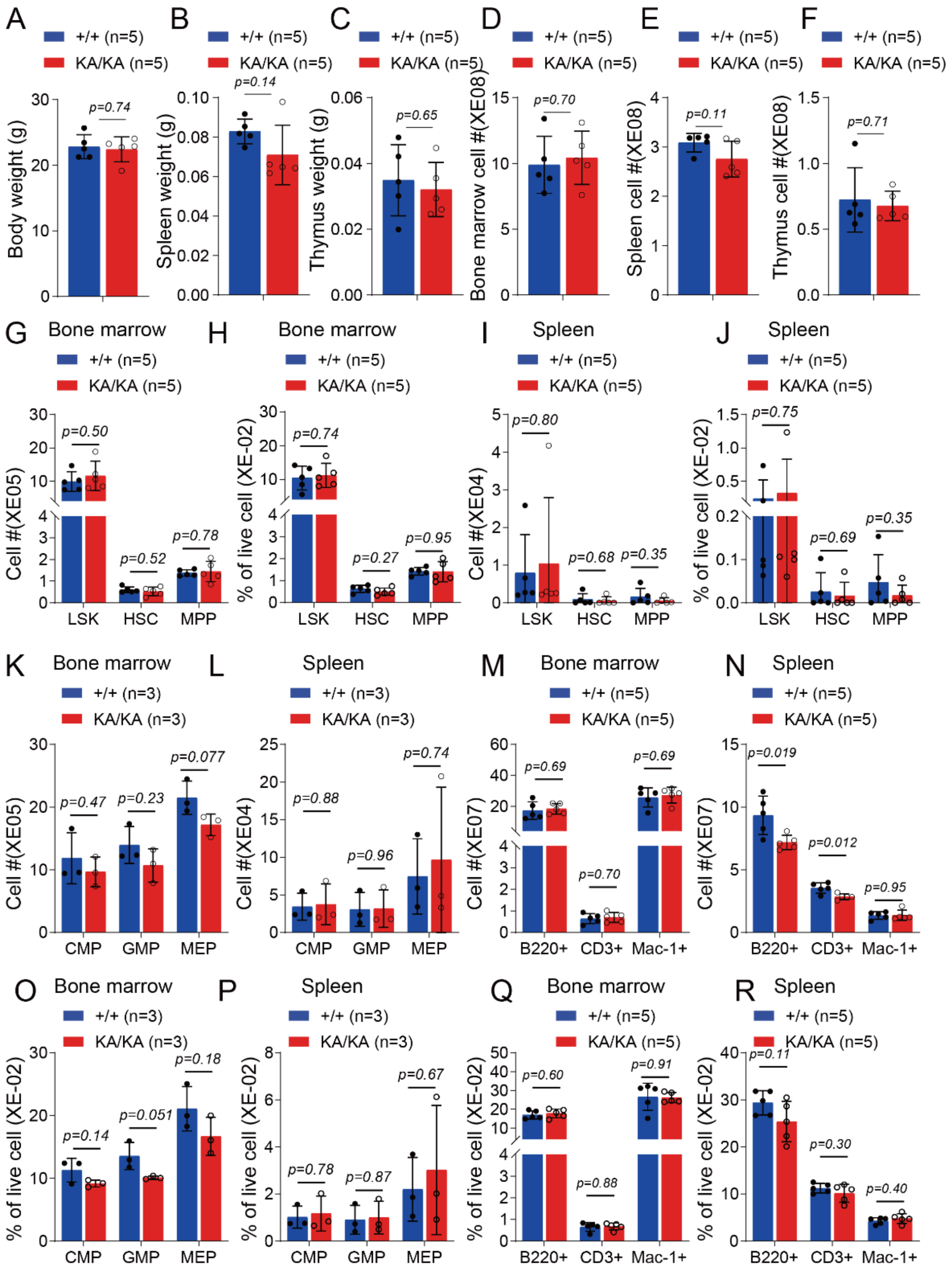


**Supplementary Figure 8. Steady state hematopoiesis in *Vav1-cre*$^+$*; PERK K618A*$^{fl/fl}$ mice.** 6 to 8 weeks old *Vav1-cre*$^+$*; PERK K618A*$^{fl/fl}$ (fl/fl) and *Vav1-cre*$^-$*; PERK K618A*$^{fl/+}$ or *Vav1-cre*$^-$*; PERK K618A*$^{fl/fl}$ (+/+) mice were analyzed for body weight **(A)**, spleen **(B)** and thymus **(C)** weight, cellularity of bone marrow **(D)**, spleen **(E)** and thymus **(F)**, frequency and numbers of HSPC **(G-J)**, numbers of lineage restricted progenitors **(K-L)** and mature blood cells **(M-N)**, frequency of lineage restricted progenitors **(O-P)** and mature blood cells **(Q-R)**. Data represent mean±s.d. Each replicate represents a single mouse from an independent experiment. Two-sided student t-test was used for statistical analysis unless specified.

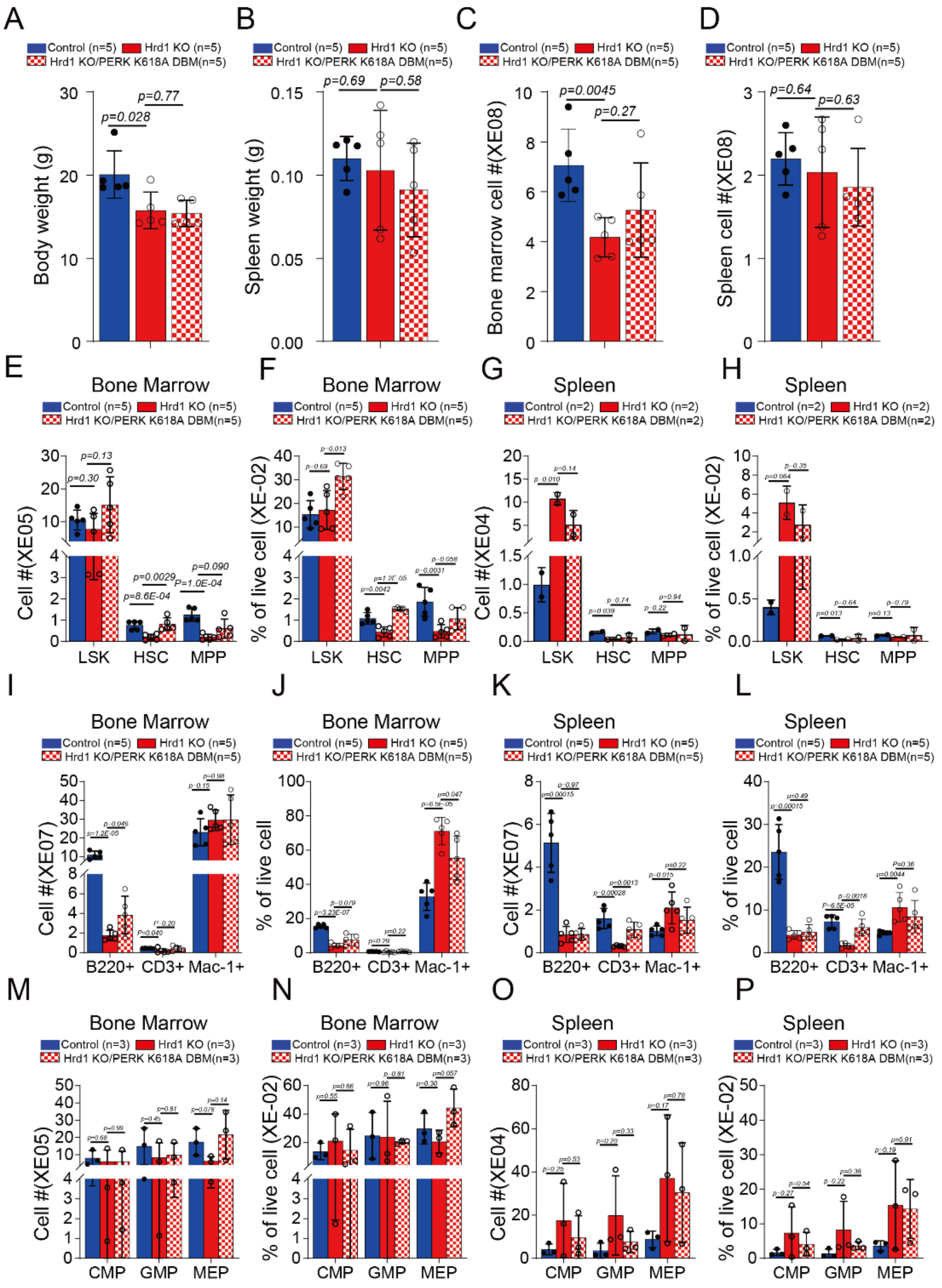


**Supplementary Figure 9 Steady state hematopoiesis in *Hrd1/PERK KA double mutant*** 6 to 8 weeks old *Vav-cre*$^{+}$*; Hrd1* $^{fl/fl}$ (fl/fl),*Vav1-cre*$^{+}$*; Hrd1* $^{fl/fl}$ (fl/fl) *;PERK K618A*$^{fl/fl}$ (fl/fl) and control (+/+) mice were analyzed for body weight **(A)**and spleen weight**(B)**, cellularity of bone marrow **(C)** and spleen **(D)**, frequency and numbers of HSPC **(E-H)**, mature blood cells **(I-L)** and lineage restricted progenitors **(M-P)** in spleen and bone marrow.. Data represent mean±s.d. Each replicate represents a single mouse from an independent experiment. Two-sided student t-test was used for statistical analysis unless specified.

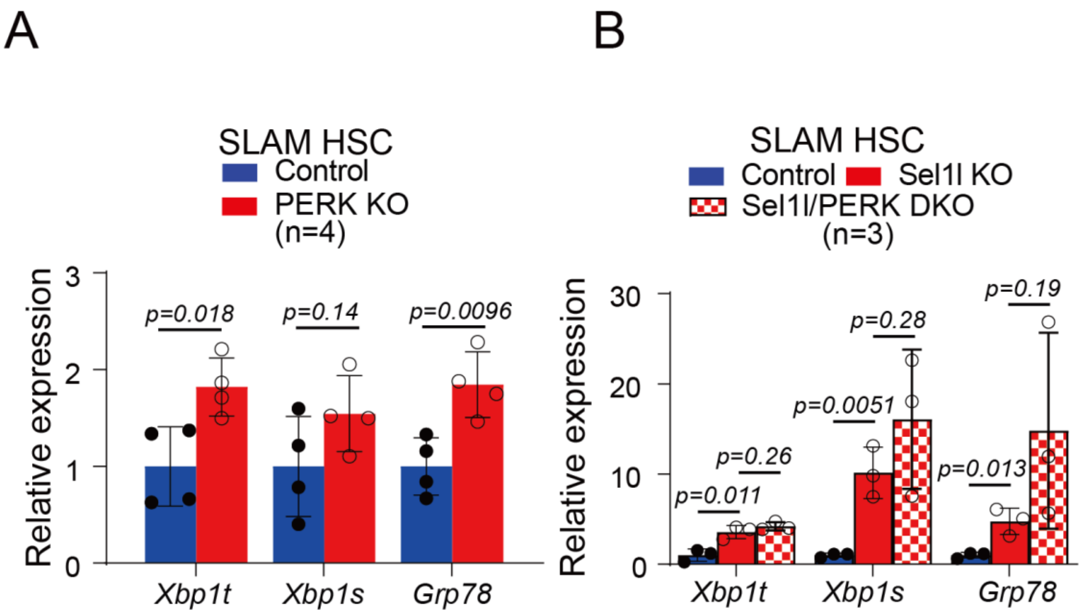


**Supplementary Figure 10 PERK depletes damaged HSCs without exacerbating ER stress**

Two weeks after pIpC injection, *Mx1-cre*$^+$*; PERK* $^{fl/fl}$ (PERK KO) and control mice (**A**), or *Mx1-cre*$^+$*; Sel1L* $^{fl/fl}$ (Sel1L KO), *Mx1-cre*$^+$*; Sel1L* $^{fl/fl}$*; PERK* $^{fl/fl}$ (DKO) or control mice (**B**) were analyzed for Xbp1s and Grp78 expression in purified SLAM HSCs (by RT-qPCR). Data represent mean±s.d. Each replicate represents a single mouse from an independent experiment.
